# A Petascale Automated Imaging Pipeline for Mapping Neuronal Circuits with High-throughput Transmission Electron Microscopy

**DOI:** 10.1101/791889

**Authors:** Wenjing Yin, Derrick Brittain, Jay Borseth, Marie E. Scott, Derric Williams, Jed Perkins, Chris Own, Matt Murfitt, Russel M. Torres, Daniel Kapner, Adam Bleckert, Daniel Castelli, David Reid, Wei-Chung Allen Lee, Brett J. Graham, Marc Takeno, Dan J. Bumbarger, Colin Farrell, R. Clay Reid, Nuno Macarico da Costa

**Author notes:** Correspondence information: Wenjing Yin, Clay Reid and Nuno Macarico da Costa. These authors contributed equally (Wenjing Yin, Derrick Brittain, Jay Borseth).

## Abstract

Serial-section electron microscopy is the method of choice for studying cellular structure and network connectivity in the brain. We have built a pipeline of parallel imaging using transmission electron automated microscopes (piTEAM) that scales this technology and enables the acquisition of petascale datasets containing local cortical microcircuits. The distributed platform is composed of multiple transmission electron microscopes that image, in parallel, different sections from the same block of tissue, all under control of a custom acquisition software (pyTEM) that implements 24/7 continuous autonomous imaging. The suitability of this architecture for large scale electron microscopy imaging was demonstrated by acquiring a volume of more than 1 mm^3^ of mouse neocortex spanning four different visual areas. Over 26,500 ultrathin tissue sections were imaged, yielding a dataset of more than 2 petabytes. Our current burst imaging rate is 500 Mpixel/s (image capture only) per microscope and net imaging rate is 100 Mpixel/s (including stage movement, image capture, quality control, and post processing). This brings the combined burst acquisition rate of the pipeline to 3 Gpixel/s and the net rate to 600 Mpixel/s with six microscopes running acquisition in parallel, which allowed imaging a cubic millimeter of mouse visual cortex at synaptic resolution in less than 6 months.

## INTRODUCTION

Serial-section electron microscopy has a long history of elucidating brain structure and connectivity^1^; more than sixty years after the first uses of electron microscopy in neuroscience^2^, it remains the sole method that permits complete morphological analysis of every neuron in a sample, along with identification of all chemical synapses. With recent technical advances, there has been a renaissance in the field. The first complete connectome, of an entire worm nervous system^3^ has been updated and extended^4,5^, while progress in system automation^6–11^ has enabled large scale studies of the nervous systems in the fly, fish, bird, and mammal^7,10,12–22^. In some cases, like the fruit fly^10^, the dataset includes the totality of the adult brain, while in mammals the largest dataset (~0.07 mm^3^) includes a fraction of a thalamic region^20^ (incidentally the same structure, the dorsal lateral geniculate nucleus, was the target of one of the largest 20^th^ century reconstructions in the mammalian brain^23^).

A longstanding goal has been to reconstruct a complete local cortical microcircuit. Depending on the species and cortical region, this might require imaging one cubic millimeter or more, at least an order of magnitude larger than previous datasets. Imaging at the scale of a cubic millimeter is best done on EM systems capable of handling thousands of sections at a time, offering fast imaging without sacrificing resolution. Such EM systems should also provide high degrees of automation and reliability in order to permit unsupervised continuous operation.

Our overarching strategy is to combine transmission electron microscopes (TEMs) with customized, precise sample handling and fast, high-resolution cameras to increase the net imaging rate^7,10,14,19^. Automation, system robustness, and integrated quality control all proved essential to collect data from a very large volume with an extremely low error rate. TEMs achieve perhaps the highest signal-to-noise ratio^10^, especially for fast imaging, but commercially available TEMs are neither designed nor optimized to efficiently image serial sections at a large scale. Most commercial TEMs can hold just a few sections at one time and assume manual operation, however for the dataset considered here more than 26,500 serial sections were collected, therefore requiring a highly automated approach. Previously, automated sample control had been achieved with a system for handling grids^10^ or with the GridTape approaches used here^24^. Because the imaging of serial sections is a highly parallelizable task, we implemented a distributed platform of multiple high-throughput automated Transmission Electron Microscopes (autoTEMs) that can image in parallel different sections from the same block of tissue. We refer to this pipeline as Parallel Imaging using Transmission Electron Automated Microscopy (piTEAM). The piTEAM infrastructure is unique as a process to sustain parallel imaging and its major novelties are that each TEM is almost completely autonomous and imaging can be coordinated over multiple microscopes. This allows for massive scaling of acquisition rate. The pipeline offers an exceptional combination of speed, resolution, signal-to-noise, and cost efficiency to map the brain structure and neural circuits. Parallelization over multiple microscopes also adds robustness to system failure and downtime, since no single microscope is a bottleneck for data production. Throughput can be increased either by upgrading components, such as cameras, in individual microscopes, or by scaling horizontally with more microscopes. Each autoTEM is a more automated version of the first generation of TEMCA (Transmission EM Camera Array) instruments^7,14,19^ where the column of the TEM was extended to enlarge the projected image on a large phosphor screen, which was then imaged by an array of cameras to achieve a large field of view (FOV) at high resolution. We have built on the original TEMCA design to create a system that has full automation, simple modular design, and systems-level feedback for error correction and quality control, allowing piTEAM to operate with little user intervention after the initial experimental setup. The acquisition software (pyTEM) has been designed in-house for continuous autonomous imaging, supporting real-time centralized imaging control across multiple systems. The piTEAM infrastructure integrates imaging automation at the level of TEM control, TEM calibration, TEM monitoring, and a tissue database which allows us to create highly standardized datasets and to be able to swap sections across microscopes without the need for manual calibration or manual region of interest (ROI) setup.

In the current platform, the burst imaging rate of each autoTEM is 500 Mpixel/s (image acquisition only) and the net continuous imaging rate is 100 Mpixel/s (image acquisition and operational overhead time such as focus and exposure adjustments, stage movement, image capture, QC, and post processing). Deploying 6 autoTEMs yields a combined burst acquisition rate of the pipeline to 3 Gpixel/s and the net rate to 600 Mpixel/s. This imaging platform has proved its reliability by imaging multiple datasets that contain the full depth of the cortex from pia to white matter, with imaging ROIs of ~1 mm^2^ per section. The imaging rate was reached while acquiring a petascale mouse neocortex volume > 1 mm^3^ of mouse neocortex that spanned four different visual areas. More than 26,500 thin slices of tissue sections were successfully imaged within six months. During this period, five microscopes imaged 24/7 with an average uptime of 65%. This is to our knowledge the longest series of sections imaged with high resolution (~4 nm pixel size) by electron microscopy. These sections were collected onto tape and imaged with a reel-to-reel system^24^.

To accomplish this challenging task, three key elements were kept in mind when designing the EM imaging pipeline: fast acquisition, high image quality, and system robustness. Great efforts were devoted to improving the system automation and reliability. Stage motion and exposure times were optimized to increase acquisition speed. Closed-loop feedback systems were created to protect sections, the stages, and the microscopes during imaging, as well as for error handling and reimaging down the road. The microscopes and hardware components also underwent several iterations of validation and design changes to provide favorable vacuum conditions and beam stability for the high-current lanthanum hexaboride (LaB6) type electron source. Crucially, images were optimized with automated procedures such as autofocus, automated flat-field correction, and real-time quality control, to ensure consistent high quality.

## RESULTS

### Development of an automated Transmission Electron Microscope (autoTEM)

The imaging platform described here uses a standard JEOL 1200EXII 120kV TEM that has been modified with customized hardware and software to increase effective imaging rate and automation. We chose this microscope, first produced in the 1980s, because the hardware is very robust and for its relative abundance in the resale market, making it a very cost-effective platform. The key hardware modifications to the stock JEOL TEM include: (1) an extended column and custom 12” electron-sensitive scintillator that produce a 10-fold increase in the field-of-view with negligible impact on spatial resolution, similar to the original TEMCA design of Bock and colleagues^7^ but with a larger scintillator; (2) a large-format CMOS camera outfitted with a low distortion lens that reduces image acquisition time to 50-150ms with low field distortion; (3) a nano-positioning sample stage that offers fast, high-fidelity raster imaging of large tissue sections; and (4) a reel-to-reel tape translation system that accurately locates each aperture using index barcodes for random access on the GridTape developed by Lee and colleagues^24^.

The column extension allows a larger image to be formed on the custom scintillator, which is imaged by a single CMOS camera in a custom housing. The camera is positioned using a stage with precision of micrometers for linear and rotational adjustment, which allows for a lens with large aperture size and consequent shallow depth of field. Field distortion is minimized using a high-quality commercial lens (Zeiss Otus, 55 mm f/1.4). Using a single large sensor rather than the original 2×2 camera array^7^ eliminated the computational overhead required to queue, process, and quality-control images from separate cameras, thus improving system robustness while maintaining high throughput. For the mouse mm^3^ dataset described below, we used initially 20 Mpixel cameras (XIMEA, CMOSIS CMV20000), some of which were later swapped for 50 Mpixel cameras (CMOSIS CMV50000) that immediately doubled throughput (burst rate: ~0.5 Gpixel/s, net rate 0.1 Gpixel/s) so that ~70 montages, each 1 mm^2^, could be imaged per day. These large-frame high-speed sensors were unavailable when the first TEMCA system was created^7^, and demonstrate how this approach takes advantage of the tremendous progress in camera sensor technologies. The use of off-the-shelf cameras decreased cost while permitting upgrades when new sensors arrive on the market.

Most standard TEMs can handle 1 to 5 sections per load, but recently systems have been designed that can handle hundreds to thousands of sections^10,24^ thereby allowing continuous imaging of serial sections without interruption. A series of custom sample stages with increasing sample packing density and microscope automation were constructed to achieve the present automated pipeline. Presently, we have implemented two different stage designs: a piezo two-axis stage with high-density standard sample grids (Voxa GridStage Sprite^™^, more details discussed in Methods) and a TEM tape-compatible stage (Voxa GridStage^™^) that can handle thousands of sections in one sample load. The tape approach to TEM was pioneered by K. Hayworth and colleagues and was newly engineered to be GridTape^24^. The tape is manufactured from Kapton which is robust but not electron-translucent, therefore and in order to make TEM-compatible, GridTape incorporates a 50-70 nm LUXFilm (Luxel) into windows that have been laser-milled into Kapton tape^24^. We have used GridTape on the mm^3^ data collection detailed bellow, using an Automated Tape Collecting Ultra-microtome (ATUM)^11^, an established approach for reliable automated section collection^17,20^.

The reel-to-reel system (Figure 1e and 1g, details in Methods) used in our pipeline allows for up to 5,500 sections on GridTape to be loaded in a single pump down cycle. The elimination of repeated sample loading allows for 24/7 automated imaging over large sample sets. Since each sample has a unique barcode ID, the system also allows random access of individual sections via spooling to the desired position. Two tape reels located on either side of the TEM column supply specimens and collect them after imaging. The vacuum load lock for inserting the GridStage sits on top of the feed reel and includes a tension sensor to automatically adjust the tape movement when changing apertures. Samples proceed through a pinch drive motor system, tension sensor, and then are fed through the channel of the GridStage. The tape is always tensioned during movement and is slack during montaging which mechanically and thermally isolates the imaging region to facilitate high quality acquisition. Clamps within the GridStage mitigate micro-vibrations and eliminate tape slippage during fast scans that can disrupt absolute position accuracy. Two additional camera viewports are also provided in the feed and take-up reel housings, which are used to monitor tape status during sample translation.

**Figure 1:**
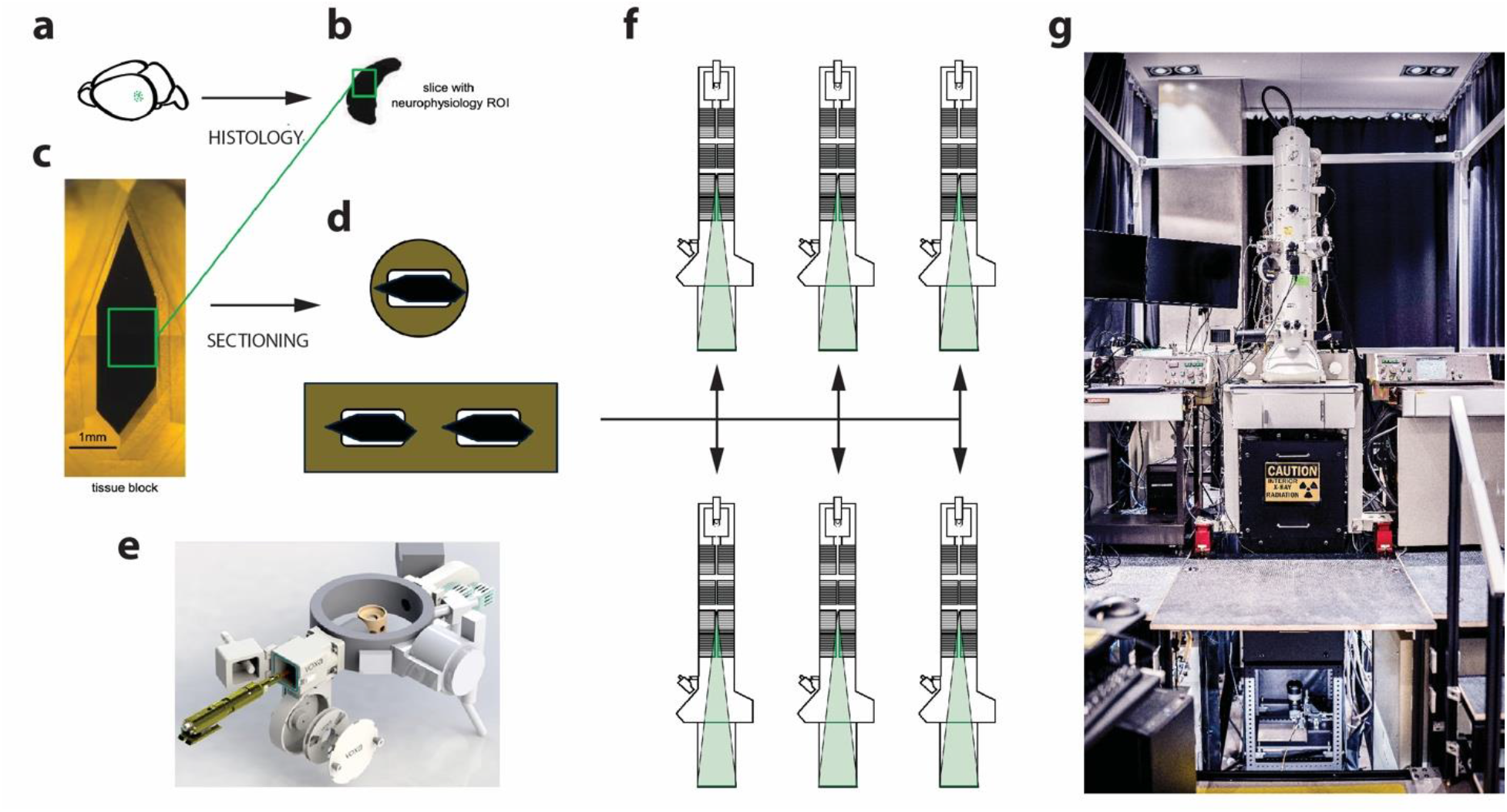
Experimental pipeline from sample preparation to piTEAM imaging. a. Mouse brain with region of interest indicated with green dots. b. Slice of brain after osmium protocol, the green square represents the region of interest (ROI) to be imaged with the electron microscope. c. Block of tissue trimmed and prepared for sectioning. d. Schematic of ultrathin sections on grid and tape. e. 3D rendering of GridStage Reel and its reel-to-reel sample translation system. f. Cross-section schematic of the distributed autoTEMs. g. Photograph of an autoTEM system in the EM suite.

In order to collect a petascale data set over months of continuous imaging, we designed the autoTEM system to be almost completely autonomous. In the sections below we describe the infrastructure that enabled this automation starting with the software. Automation was a crucial step for scaling and to replicate this autoTEM system over six microscopes, allowing simultaneous imaging of multiple tapes.

### A software infrastructure for petascale imaging via parallel imaging transmission electron automated microscopy (piTEAM)

An automated electron microscopy pipeline such as piTEAM requires data-driven, systems level control similar in principle to the *fly-by-wire* approach to automation in avionics to digitally control flight systems using a closed feedback loop. A similar principle is used by the autoTEM system to control the state of the microscope without human intervention, dynamically changing parameters to ensure consistent data quality and throughput. Live measurements are collected (e.g., focus score range, pixel intensity histogram spread, brightness uniformity, beam centering, lens distortion within FOV) and the generated data are used to adjust microscope parameters to stay within given limits defined prior to imaging. The software was designed to abstract imaging and system operation as much as possible and to support different sample handling implementations and their corresponding features as they were developed.

In summary, the image acquisition software has been designed with the following goals in mind:

- Completely autonomous operation. An autoTEM system should be able to run without user intervention beyond initializing the experiment.
- Adaptable to advances in sensor technology, enabling fast and affordable scaling. We have developed an abstracted software architecture, functional validation test plan, and change management plan to understand the impacts of sensor upgrades on image quality and effective throughput.
- Automatic ROI generation from optical images or low-mag EM images.
- Remote control of the TEM machines using web technologies.
- A cloud-based database, associating specimens, tapes, apertures, and ROIs which span optical and EM imaging modalities. This database manages the millions of images that are generated and tracks them back to ROIs, section ID, microscope configuration, imaging condition, etc.
- Remote multi-system monitoring (MSM) tool that analyzes system and environmental variables (such as temperature and vacuum, among others) and uses feedback mechanisms to prevent sample or hardware damage.

The core of the image acquisition software is divided into three components that work in synchrony to provide continuous uninterrupted acquisition (Figure 2a and 2b). The first component is the *pyTEM* software package which is a set of Python modules that provide orchestration and control of the EM hardware, stage, and tape sub-systems. This core control software is built upon a state machine which accepts high-level commands from the *pyTEM GUI* (Figure 2c and S2-4). pyTEM controls the microscope via serial ports, calculates the positions of the images to be captured, issues the commands to the stage and reel-to-reel systems, writes metadata files, sends system status messages to multi-system-monitoring (MSM), and publishes preview images and status to the pyTEM GUI. Time sensitive operations are processed by the GPU-based *TEM Graph (*described below), whereas less time critical functions such as auto exposure and auto focus are integrated into pyTEM.

**Figure 2.**
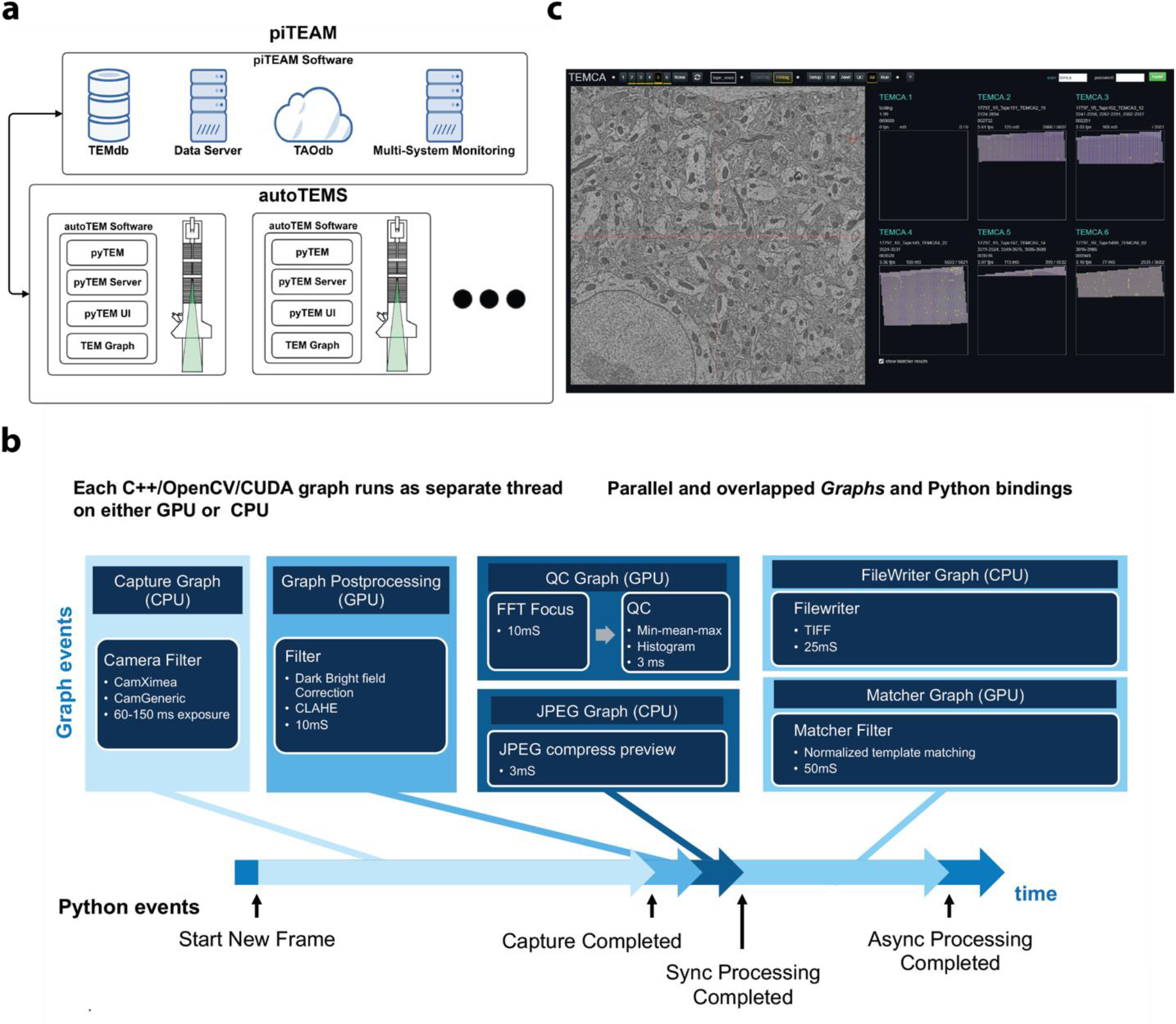
a. The architecture of piTEAM pipeline. The aperture ROIs (see figure 1d) are uploaded to AWS and are pulled from TAO DB for imaging. A pyTEM server allows for remote control and the MSM monitors system and environmental health. b. TEM Graph key components. Images are acquired and loaded into GPU memory. A series of filter graphs apply corrections to the image (flatfield, down sampling for GUI preview). Separate graphs check image quality and statistics while the image is written to disk in parallel. c. pyTEM GUI. The left EM image is a preview while the right is an example of parallel imaging on five systems for 1 ×1 mm^2^ montage. The pyTEM GUI provides the user with an intuitive, web-based interface to perform manual imaging surveys as well as long serial montage runs containing hundreds or thousands of ROIs. From the web-UI any running autoTEM system can be observed and controlled.

The second component is the TEM Graph which is a group of custom filter graphs allowing real time image processing. To achieve the highest possible acquisition rates the lowest level component in the system, TEM Graph is written in C++ using CUDA for GPU processing to maximize acquisition rates. The design is also highly parallel, each graph component runs on a separate thread, and certain graph components can run asynchronously. TEM Graph includes Python bindings for control and event notification to pyTEM, which provides high level control of the entire acquisition system. The TEM Graph allows for parallel, asynchronous operations and multi-GPU configurations, some examples of its features include (1) image correction (flatfield image correction, contrast limited adaptive histogram equalization (CLAHE) enhancement); (2) feature/point matching (template matching); (3) real-time image quality metrics (autofocus, histogram, standard deviation).

The third component *pyTEM Server* interfaces with many systems in parallel. It was designed to allow control of any autoTEM system with multi-user access remotely. A single server is dedicated to a single system. The server hosts the pyTEM GUI web pages and provides a link between pyTEM and multiple web clients.

Additionally, two high-level modules complete the acquisition software: the pyTEM GUI and the Multi-System Monitoring (MSM). The pyTEM GUI (Figure 2c) is a web-based interface to perform manual imaging and serial montages. A single master web interface can switch dynamically to control any TEM in the pipeline and support multiple simultaneous GUI clients. pyTEM GUI is also used for ROI definition (Figure S3), monitoring of automatic optical ROI QC, system calibration, monitoring, and montage initiation. pyTEM GUI uses responsive web design principles and operates at any screen resolution even on a mobile phone (Figure S4). MSM (see also supplemental information, Figure S6 and S7) is a combination of a Python web server and web-based UI that allows the users to quickly ascertain if any subcomponent needs attention. Time series data are recorded to a database, and if a system is detected to have an issue, MSM communicates with pyTEM and informs the appropriate users that an error has occurred. In catastrophic failure modes, MSM can also pause the acquisition of an autoTEM system to prevent system damage and potential sample loss. An example of such failure mode that we have experienced is a break of the water-cooling system, where the MSM stopped acquisition and safely returned the scope to low mag to reduce potential sample damage from excessive beam exposure.

During serial sectioning, database entries are automatically created for each indexed sample aperture in the Amazon Web Services (AWS) cloud. Each aperture is represented by a *TEM Acquisition Object (TAO)* (see supplemental information, Figure S8) that includes the ROI to be imaged, an optical image acquired during sectioning, the index barcode from this image, and information about the specimen and substrate (Figures 1 and 2). ROIs can be defined automatically during upload to the *TAO DB*, as a separate machine vision task running either locally or on AWS, or manually via the pyTEM GUI. Once ROIs have been defined for a region of GridTape, imaging can commence.

When montaging, pyTEM GUI first receives the initial user input such as random aperture ranges and then the entire acquisition by pyTEM and TEM Graph is autonomous. The system will continuously loop through the following steps (Figure S5): 1) advancing the tape, 2) seeking barcode, retrieving the corresponding TAO, 3) extracting the ROI and 4) montaging the assigned aperture until it either completes acquiring the desired range of ROIs or an error event is detected to trigger an abort.

A few montage preparation steps are also automatically initiated for each ROI, such as centroid finding, flatfield correction, and autofocus routine. If the flatfield and autofocus return values within preset thresholds, the montage raster scan starts. TAO DB can be accessed by multiple machines simultaneously for parallel imaging. Each acquired image is displayed in pyTEM GUI along with real time QC statistics and tile overlap measurements.

### Integration of fully automated closed-loop feedback and real-time QC

Beyond the difficulty of automating TEMs, a further requirement for the imaging pipeline is to ensure that the multiple, concurrently active systems are consistently producing high quality data and preserving sample integrity. The system is designed with a fully automated closed-loop system that can provide robust error handling and enable the machine to execute appropriate responses based on simple quality trigger metrics. Figure 3 shows the data flow through the piTEAM pipeline. The digitally controlled microscopes receive acquisition parameter inputs from a user and extract ROIs from AWS cloud database. Before each montage starts, MSM checks the environmental health conditions such as filament, vacuum, water pressure, and temperature for lens cooling etc., and decides whether to proceed with imaging or to stop the acquisition and put the system in a safe condition. Beam position and focus are also verified before montaging. If the beam is displaced to a corner of the FOV, an “automated beam centering” routine will be launched to reposition the beam to the center of the screen. Deficient brightfield or focus show as alerts on the montage status feedback and the system automatically reattempts to re-acquire the montages. Real-time QC is performed during montage, collecting image quality information such as focus score, tile overlap error, and image histogram into a metadata file along with the raw acquired images. Once a single montage is completed, the system automatically analyzes the QC information and makes a risk assessment on whether to proceed to imaging the next indexed sample. If the first-pass QC result is good, the data will be sent on to the storage server, otherwise it is marked as a QC failure. If the QC failures exceed a user-set threshold, the acquisition automatically stops.

**Figure 3:**
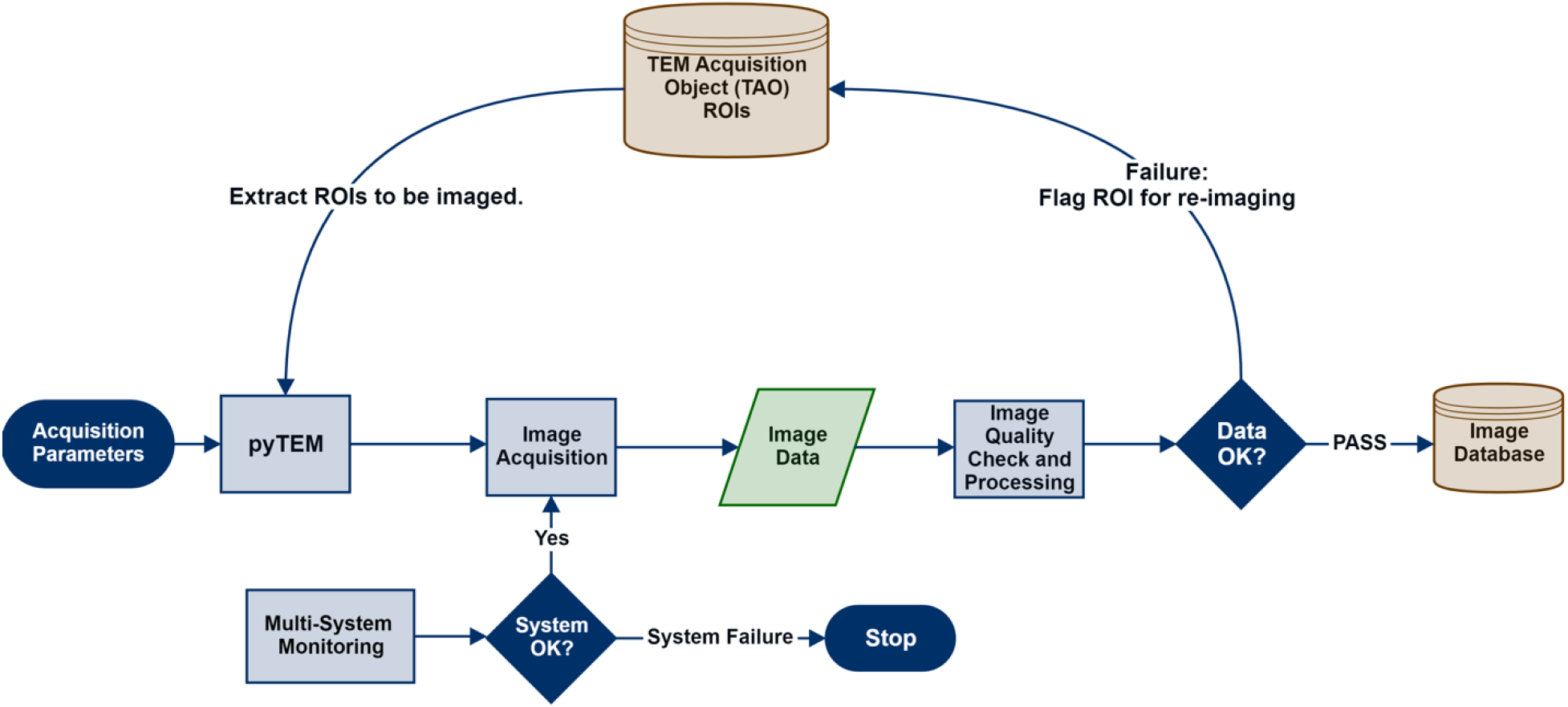
Closed-loop image data flow. After pyTEM receives ROI’s and acquisition parameters, image acquisition is triggered and image data is then analyzed on-the-fly on the acquisition computer. Rejected image data (those failing to meet QC thresholds) are flagged in a pyTEM DB instance to be re-imaged. The database will be updated to reflect which ROIs need reimaging. If the montage passes inspection, it is sent to a data center for post-processing, alignment and storage.

The QC module was tested during the collection of a dataset containing tens of thousands of 2D montages over 100 million individual tiles. The collection of such a vast dataset allowed us to encounter and correct various imaging errors that called for reimaging of sections. Detailed examples and images are discussed in supplement S11. Such errors are best corrected during imaging because they might cause long delays on image processing and data analysis later, therefore a need for real-time montage validation during acquisition becomes necessary. To ascertain whether the data generated are of sufficient quality for post-processing, a real-time image quality control software package has been developed.

Real-time QC is based on GPU processing of tile matching and embedded in the pyTEM imaging software. This module captures image and system errors in real-time and displays the errors on pyTEM GUI. The matcher filter utilizes normalized cross correlation template matching at tile edges to check the overlap between neighboring tiles along both X and Y directions (Figure 4a). Tile overlap measurement is carried out during acquisition and results are saved in metadata file for each individual montage. The fast GPU processing allows immediate visualization and identification of imaging problems and triggers reimaging without impacting the rest of the pipeline running on CPU and system memory. Typical average performance for three overlap matches along the top and three matches along a side edge is ~50 ms, a fraction of an acquisition frame. Real-time QC overlaps with the next stage move in time, and sustainably avoids extra total acquisition overhead.

**Figure 4.**
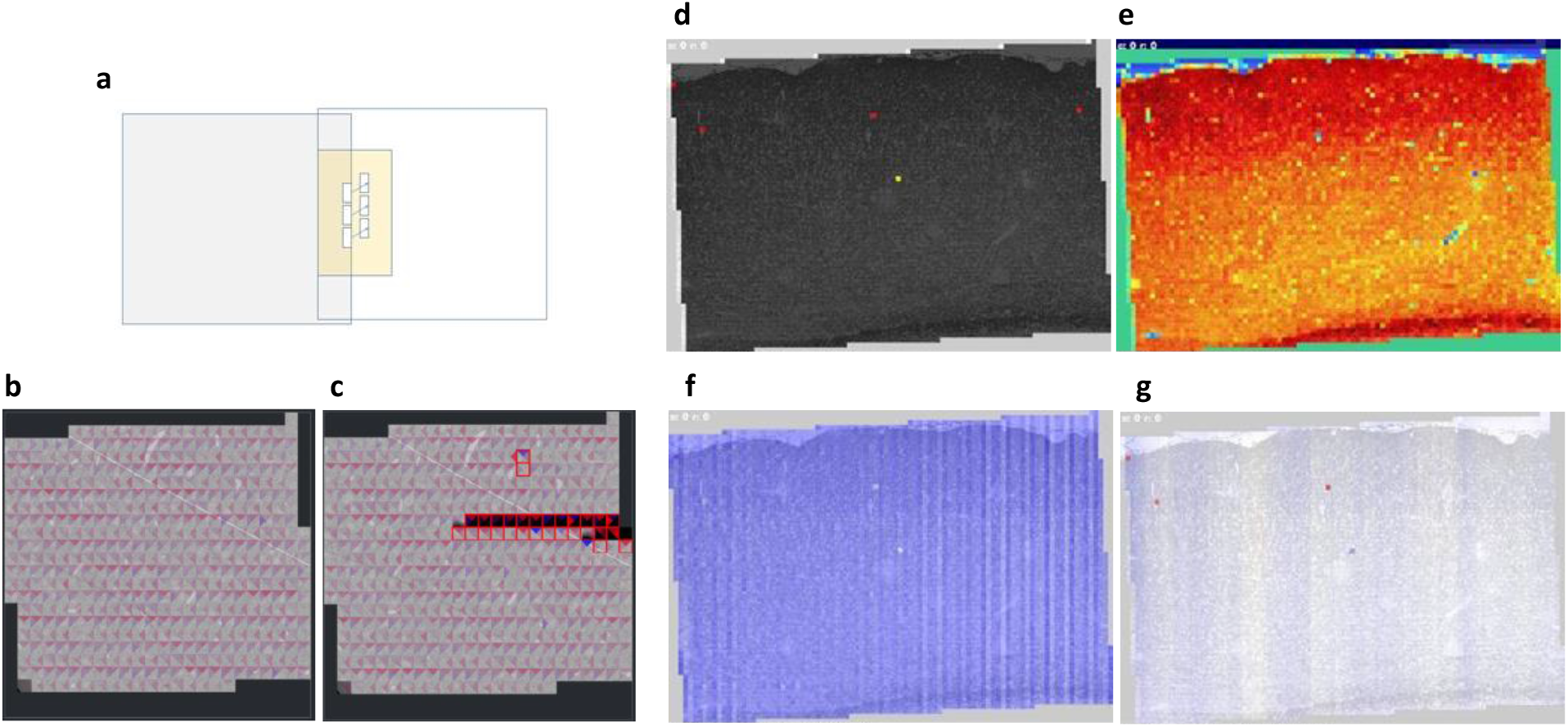
Real-time quality control (QC) for capturing image and system errors. It utilizes the template matching to detect any tile overlap issue during acquisition, and FFT score to measure the focus quality of each tile. a. Source templates are 5% of image side. This ensures that a minimum of 5% image overlap exists for the downstream pipeline. Three templates are used per edge, and the mean and standard deviation of the three matching vectors are returned from the filter. In general, the standard deviations between the match vectors is less than one pixel, which indicates that the match accuracy is excellent. The template search area is a region of twice the tile overlap (~ 13% for 20MP camera). b. Good vs. c. bad real-time matcher results displayed on GUI: each triangle represents a match result, which the color represents the angular direction between the ideal position and the template matched position, and the intensity represents the magnitude of the offset between the ideal and actual locations. The right bad montage flags the errors of misaligned rows and dark tiles. Imaging problems are usually detected as flagged tiles in the quality map or non-uniform output maps. d. matcher quality map; e. focus map; f. x-offset from ideal; g. y-offset from ideal.

Figure 4b shows a typical visualization of the results of the matching operation. Blue hues indicate that the template was found beyond the expected position, and red hues indicate that the template was found before the expected position. The intensity indicates the magnitude of the deviation from the expected position. It is important to note that the matching succeeds even in areas without tissue. To test the matching operation, we introduced two types of errors shown in Figure 4c. The stage position was artificially perturbed on row 4, and then on rows 10-11 the beam was blocked, resulting in black tiles. If the number of matched templates or the standard deviation of the template match vectors fail to meet thresholds, the tile is marked as an error, highlighted as a red rectangular box. If lens calibration is performed prior to the matching operation, it can also directly produce a 2D montage from matcher results alone.

In addition, a suite of batch processing Python scripts can run through all the montage tiles offline and quickly review montage quality before sending them to image processing. The script can capture different errors through the color maps of “matching overlap quality”, image focus, image offset in both “x” and “y” dimensions. Figures 4d-g show the QC output from a good example of 2D montage. The uniform color pattern across the frame for match quality (Fig. 4d) and match distance (Fig. 4f and 4g) indicate good overlap area in-between neighboring tiles. Figure 4e shows a fairly uniform focus map, with tissue structures such as blood vessels being highlighted. Overall, this real-time QC module enables early identification of imaging errors and efficiently performs montage reimaging wherever necessary with minimal user intervention.

### Petascale data acquisition using piTEAM

The first dataset collected by the piTEAM system defined above contained 2500 tissue sections of 240 μm ×150 μm in area size. The voxel resolution for neuroanatomical dataset was 4 nm × 4 nm × 40 nm with the z dimension (40 nm) set by the sectioning thickness on the ultramicrotome. Running by 20MP XIMEA camera and a Sprite Stage sample holder (details in Supplement, Figure S1), the dataset was completed in one week using a single autoTEM system. It contained about 500 neurons and 3.2 million synapses.

The biggest challenge for scaling from this 0.03 mm^3^ to 1 mm^3^ (300× increase) was throughput. We expanded the imaging platform from a single microscope to a multi-scope pipeline parallel both in terms of sensors and electron sources, which allowed us to scale the imaging system while maintaining the large electron doses required for fast exposure. We also transitioned from a low capacity (10s of sections) stick-type Sprite Stage to a high capacity (1000s of sections) reel-to-reel with GridStage system for translating GridTape^24^ between sections and then montaging. Over 26,500 sections were divided onto seven reels that were continuously imaged across five microscopes for almost six months. With the 20MP camera, the net image acquisition speed (imaging, stage step-and-settle time, imaging overhead and image correction) was 3.3 frames per second (fps) on average. Each frame contained 3840 × 3840 pixels. Each 1 mm^2^ montage was composed of roughly 6,000 15 μm × 15 μm tiles (Figure 5c) with an overlap of 13% between tiles in both X and Y directions. The 13% was chosen in order to ensure that stage imprecisions will still allow a 7% overlap required in order to provide enough feature matches for stitching. The total file size of a single montage was about 80 GB and resulted in a daily throughput of 3.6 TB per system for continuous imaging.

**Figure 5:**
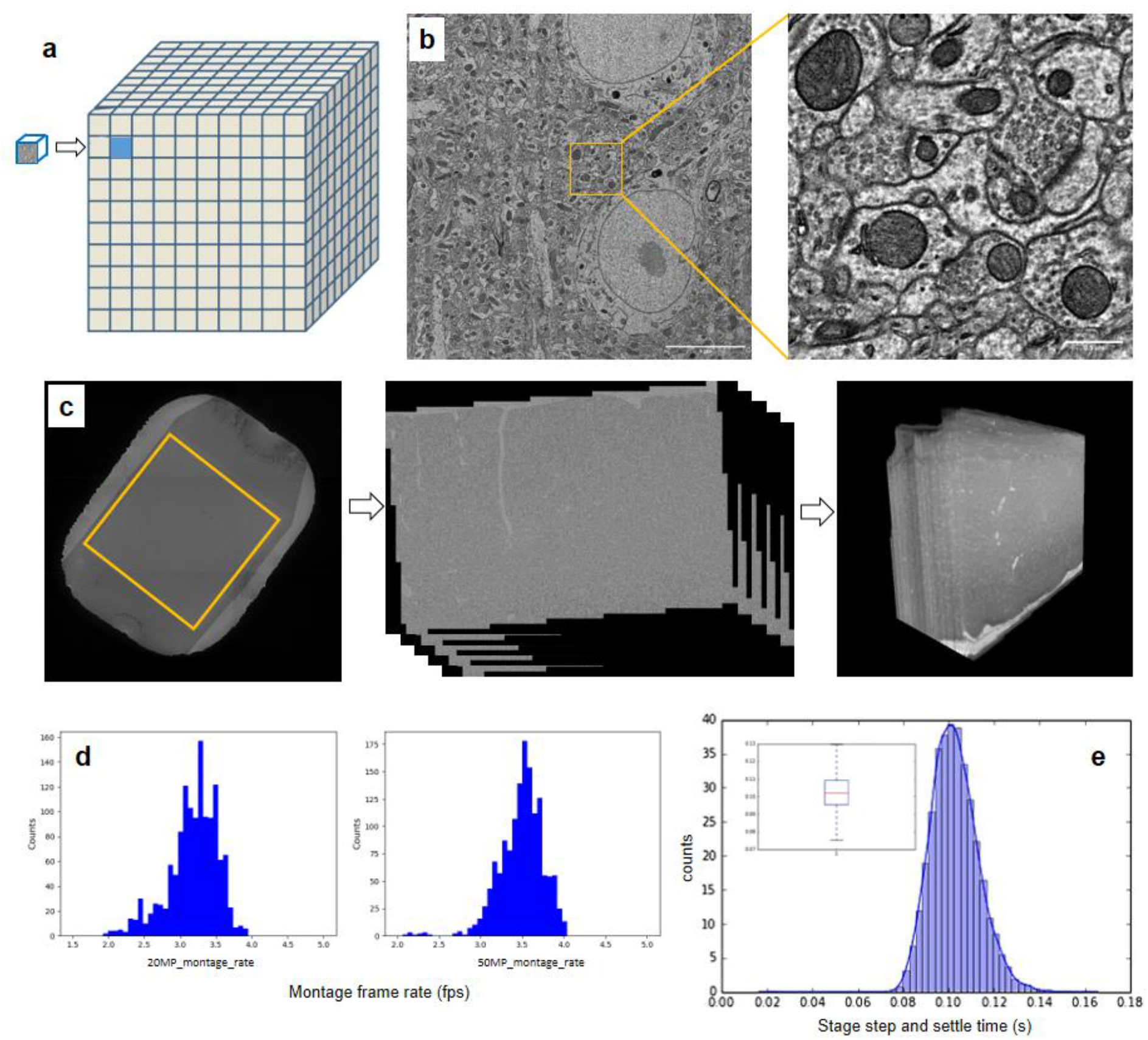
Imaging of a cubic millimeter of mouse cortex with piTEAM a. Scaling from (100 μm) ^3^ to 1 mm^3^, a 1000-fold volume increase; b. High-resolution electron microscopy image from the 1 mm^3^ dataset (scale bar 5 μm), and a zoomed-in area at synaptic resolution (scalebar 0.5 μm); c. Low-mag EM image of an aperture; 2D stacked montage minimap and aligned 3D volume. d. Distribution of montage acquisition rate (frames per second) achieved during 1 mm^3^ production. The plots represent a sample size of over 1000 sections imaged by 20 Mpixel and 50 Mpixel cameras each. The average imaging rate is between 3.2~3.5 fps. e. Example of stage step-and-settle time distribution for GridStage, averaged near 100 ms.

During production, three autoTEMs were upgraded to a 50MP camera, which increased the frame size to 5408 × 5408 pixels (21 μm × 21 μm). Figure 5b shows a high-res 50MP tile and a zoomed-in area at synapse level. The total number of tiles required per montage was reduced to ~2,600 from ~6,000 at an overlap of 9% in both X and Y, which maintained the same frame rate during montaging. (See Fig. 5d for montage rate distribution.) Figure 5e shows that the average stage step-and-settle time was achieved around 100ms. Table 1 is a comparison of the performance between 20MP and 50MP camera configurations in which we assume imaging a 1 × 1 × 1 mm volume. The reimaging rate for the whole dataset is roughly 10%, of which over 90% were caught by real-time QC embedded in the piTEAM pipeline and the remainder during post-processing or segmentation. The reimaging based on real-time QC output was also completed during the 6-month data collection period.

**Table 1:** Performance metric during 1 mm^3^ production for 20 MP and 50 MP cameras.

| Imaging Parameters | Avg.<br>Production<br>20Mpixel | Avg.<br>Production<br>50Mpixel | 50Mpixel<br>peak | Unit |
| --- | --- | --- | --- | --- |
| Frame Size (pix) | 3840 | 5408 | 5504 | aspect ratio of 1:1 |
| Resolution | 3.95 ~ 4 | 3.95 ~ 4 | 4 | nm/pixel |
| Tile Overlap | 13 | 9 | 9 | % |
| Net Imaging Rate | 3.2 ~ 3.5 | 3.2 ~ 3.5 | 4.0 | Hz |
| FOV per side | 15 | 21 | 22 | $\mu\text{m}$ |
| AOV | 225 | 468 | 484 | $\mu\text{m}^2$ |
| Pixels per image | 14.7 | 29.2 | 30.3 | Mpixels |
| Time per section | 30 ~ 40 | 15 ~ 25 | ~14 | minutes |
| Tiles w/ overlap | 5100 | 2600 | 2600 | per section |
| Sections per week (24/7) | ~300 | ~490 | ~700 | Sections per week |
| Time to image volume with one scope @65% uptime | 900 | 525 | 450 | days |
| Time to image volume with 5 scopes @65% uptime | 180 | 105 | 90 | days |
| Completion of project in months @65% w/ 5 scopes | ~6 | ~3.5 | ~3 | months |

## DISCUSSION

The design behind the piTEAM imaging acquisition pipeline discussed here is focused on robustness for the imaging of large datasets, imaging speed, scalability, and flexibility to keep up with continuously improving state-of-the-art digital cameras. Since we developed this scalable platform for multiple microscopes to image the same sample of tissue in parallel, we also paid attention to the affordability of any single machine, which was a primary concern about the original TEMCA systems^7,14,19^. Based on our design, if there is already an available TEM, a system that can achieve net imaging rates of at least 100 Mpixel/s can be built with additional components that cost roughly $125,000. If a microscope is not available, refurbished systems (such as a JEOL 1200EX-II) can be purchased for ~$125,000, thus making a single autoTEM more affordable than other options for high-throughput EM.

A great advantage of the autoTEM approach is the easy upgradeability. Our current piTEAM pipeline uses 50 MPixel CMOS sensors (Fig. 6) that provide a large FOV per tile. The transition from 20 Mpixel cameras reduced the collection time for each 1 mm^2^ montage from 40 min to 15 min and reduced the total number of images from 6000 to 2600 per montage. We have simulated pyTEM montage acquisition with the upcoming 100+ Mpixel CMOS sensors from major camera manufacturers (Fig. 6c) and estimated that the current imaging software and computing hardware could achieve a further 3x increase in speed (projected imaging metric in supplement Table S13).

**Figure 6:**
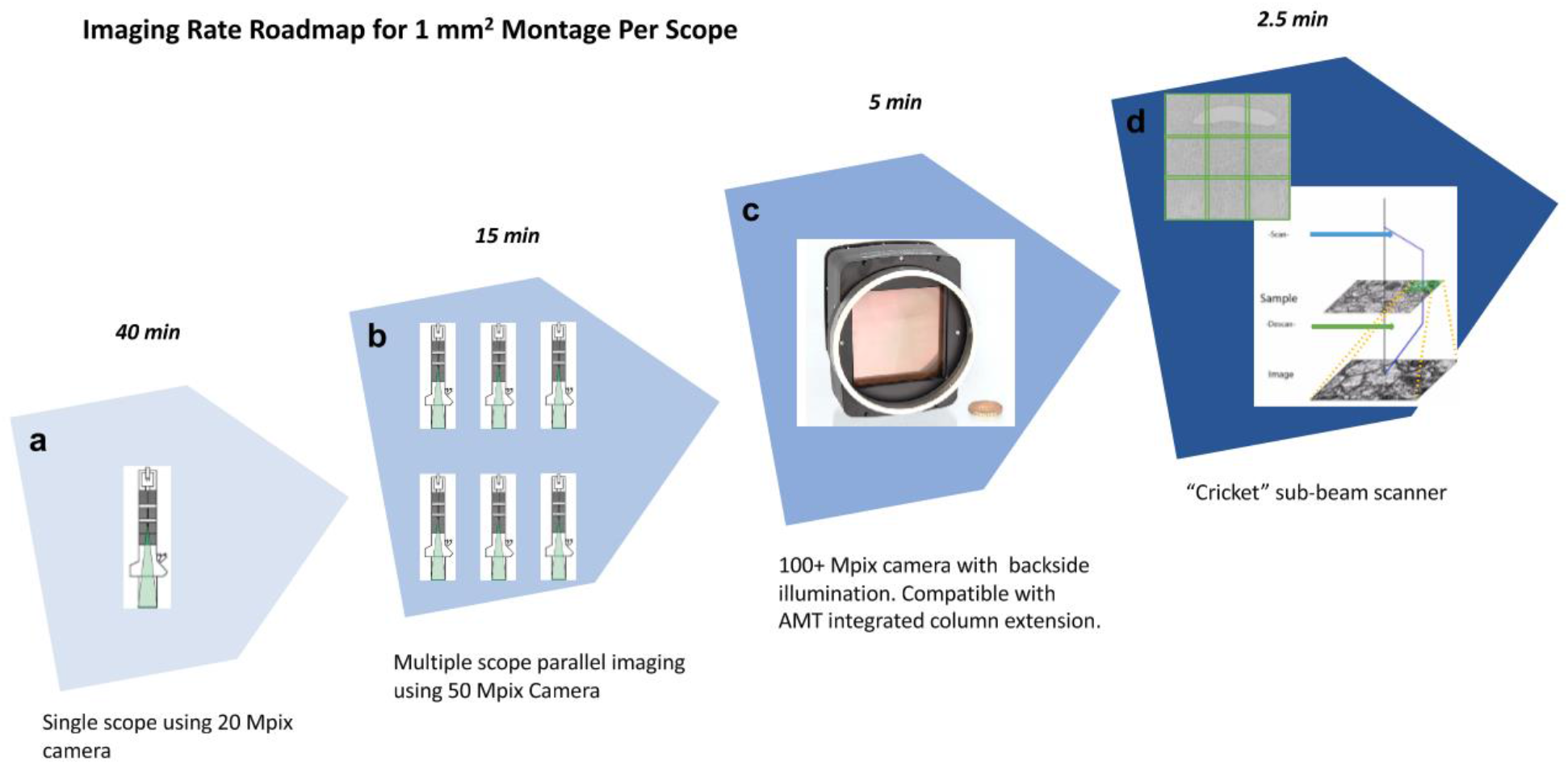
Imaging rate scaling roadmap: a. 1^st^ generation platform using 20 Mpixel XIMEA camera; b. 2^nd^ generation platform with 50 Mpixel XIMEA camera (currently the platform contains six systems for parallel imaging); c. Drop-in replacement of cameras with even larger sensor size, backside illumination and compatible AMT integrated column extension; d. An example of Cricket which is a synchronous image sub-scanner and beam blanker. The beam is scanned electromagnetically across the sample without stage movement, providing a wider field of view, and decreased overhead. The inset is a 3 × 3 supertile by Cricket beam scanner of an entire GridTape slot with tissue. The electron beam is raster scanned to allow large fields of view to be quickly imaged. The VOXA software stitches the 3 × 3 array as its output. This module can potentially be tuned for even larger FOV: 4 × 4 array.

Also, important to note is that the performance described above was evaluated on a production setting where roughly one hundred million images were collected. We have also tested several upgrades on a research and development setting that provide either a more compact design or further increases in imaging speed and scalability. As a comparison with other state-of-the-art high-throughput EM techniques, multi-beam SEM has been reported to achieve high throughput with burst imaging rates at 0.46 Gpixel/s in mouse brain with a block face approach^25,26^. This published rate however does not include all factors that lead to long-term effectives rates, such as stage motion, focus, and processing overhead. FIB-SEM^8^ was also reported to automatically generate images with superior z-axis resolution, but it does require much more custom engineering and more custom facilities and thus a much greater cost per voxel.

One of the innovations of the first TEMCA systems^7^ was an extended vacuum column, used to enlarge the final projected image. This increases the total height of the electron microscope to 15 feet, which is impractical for most facilities. We have experimented with an AMT ActiveVu lens assembly (CB500M-AV) that achieves the same magnification without the requirements to extend the height of the TEM. The lens we tested uses the XIMEA camera described above, so it was easy to integrate with our software (Figure S12). In addition, the AMT integrated lens system is also adaptable to future larger sensors to meet the requirement for scaling and serves as a good choice for the next generation column extension. Both new cameras and AMT integrated lens systems are drop-in replacements for our current autoTEMs.

Step-and-settle time of a sample stage dominates a large percentage of total image acquisition speed as the exposure time of the imaging system is reduced. The GridStage running in autoTEM consumes over 100 ms for a move to complete and trigger pyTEM to initiate camera exposure. Such stage latency issue can be mitigated by using a hybrid mode that couples stage scanning with beam scanning and takes advantage of fast deflection of the electron beam, whose response times are measured in nanoseconds. We have tested a novel beam-deflecting device (Voxa’s Inc.’s Cricket) that allows imaging multiple fields of view without moving a sample stage (Figure 6d). This beam sub-scanning system deflects the electron beam around the sample in a matrix pattern, while scanning coils below the object plane de-scan the image onto the TEM camera. The sub-tiles obtained using this “supertiling” technique are acquired and stitched together to form a large composite image, in effect creating a “virtual” camera of larger size through electron-optics and computation. A 16 Mpixel image acquired at one position can, for example, be expanded to 120 Mpixel at that same stage position using a 3 × 3 “supertile”, thus eliminating 8 stage movements. By using the sub-scanner, the platform can handle both fast high-magnification montaging and low-magnification surveying of the sample.

Finally, we are also working on a suite of software modules to perform real-time lens correction and 2D montaging using template matching. This allows us to perform the 2D stitching of complete sections “on-the-fly” during acquisition using the graphics card of the acquisition computer and significantly save computation storage and time to do the image processing afterwards. We can also monitor the change in lens distortion over time and receive immediate quantitative feedback on the quality of images and stitched visualization. We are in the process of collecting a new volume dataset and comparing the performance with the existing post-imaging feature matching pipeline.

### Conclusion

EM connectomics has seen remarkable advances in the last 10 years, making it poised to examine synaptic connectivity of neuronal networks at a very large scale. For this to happen, we took an industrialized approach to build an image acquisition pipeline, piTEAM. The entire EM imaging pipeline from sample transfer, image acquisition, and image QC is a continuous automated process. Constant feedback from all stages of the pipeline ensures data integrity and quality is maximized with minimal need for manual intervention. The transition from a prototype to an industrialized production pipeline was the most challenging problem. To facilitate this effort, we have put a strong emphasis on standardization and consistency at every step of the piTEAM pipeline, resulting in a suite of methods that are open and affordable.

The piTEAM approach allows for further increases in speed, both per microscope—by using increasingly large and fast cameras, or with beam deflection—and per facility, by increasing the number of microscopes. In 2018, we used our six-microscope piTEAM platform to collect 2 PB of EM images of 1 mm^3^ mouse visual cortex at synaptic resolution over the course of six months. We anticipate that net average rate of a single microscope to increase from the 100 Mpixel/s to 500 Mpixel/s, through a combination of larger sensors and the beam-scanning. At this rate, a single microscope should be able to image a cubic millimeter in roughly 100 days. This throughput at the cubic millimeter range make piTEAM ideal to investigate microcircuits across species, cortical regions and in health and disease, in a framework that focuses on production of brain maps at large scale. Although this pipeline was designed for connectomics, any other field requiring automated serial section imaging at the ultrastructural level can take advantage of the automation, high throughput, and affordability of the methods described in this manuscript. Our approach to TEM imaging can be used either as a single autoTEM or as a full piTEAM pipeline for distributed imaging that can be implemented both within a large dedicated facility like our own as well as distributed over a community of individual laboratories.

## Acknowledgments

We thank our project manager Shelby Suckow for her exceptional work on managing the collaboration and for keeping us aligned and on time. We thank David G. C. Hildebrand, Aaron Kuan, Jasper Maniates-Selvin and Logan Thomas for developing, milling, advice and assistance on the use of GridTape; Cliff Slaughterbeck for engineering project Management and engineering design review. T. Ayers, R. Smith and Luxel Corporation for coating tape; JoAnn Buchanan for histology; Gayathri Mahalingam for stitching and alignment; Joe Mancuso and Adam Manganiello for providing the AMT extended column and assisting experimental data collection; Davi D. Bock for his advice on TEMCA column extension and his thoughtful comments on the manuscript; Agnes Bodor for feedback on image quality; Lawrence Own and Teddy Derego for their development and continuous support in GridStage and GridCon software; and Sebastian Seung and Adrian Wanner for discussion on imaging strategies and improvements. The datasets described above were previously imaged with 2P-calcium imaging by the lab of Andreas Tolias and subsequently fine aligned and segmented by the lab of Sebastian Seung. This work was supported by the Intelligence Advanced Research Projects Activity (IARPA) of the Department of Interior/Interior Business Center (DoI/IBC) through contract number D16PC00004; NIH Grant R21NS085320 (W-C.A.L); and by the Allen Institute of Brain Science. The U.S. Government is authorized to reproduce and distribute reprints for Governmental purposes notwithstanding any copyright annotation thereon.

The views and conclusions contained herein are those of the authors and should not be interpreted as representing the official policies or endorsements, either expressed or implied, of the funding sources including IARPA, DoI/IBC, or the U.S. Government. We thank the Allen Institute’s founder, Paul G. Allen for his vision, encouragement and support.

## Author Contributions

N. M. C. and R. C. R. conceptualized the project. W. Y. conducted process integration, experiments and system validation for the imaging pipeline. D. B. and M. S. designed and built the autoTEM and closed-loop system. J. B. and D. B. designed, built and developed pyTEM software for image acquisition. J. B., W. Y. and D. B. developed real-time QC. W. Y., D. B., D. J. B., M. T. and N. M. C. performed imaging and troubleshooting of 1 mm^3^ dataset for mouse visual cortex. R. M. T. was responsible for data transfer and storage. D. B., D. W., J. P. and A. B. developed the prototype pyTEM image acquisition software. C. O. and M. M. built and developed the reel-to-reel GridStage and GridCon software. D. K. and J. B. developed real-time lens correction and 2D montage. D. C., D. R. and C. F. provided manufacturing process engineering support. W-C. A. L. and B. J. G. developed and provided a prototype reel-to-reel TEM imaging stage and tape handling machine that was used during the development stages of the pipeline, they also provided materials, advice and assistance for the use of GridTape. W. Y., D. B., J. B., C. R and N. M. C. wrote the manuscript with input from all authors.

## Methods

### VOXA GridStage Sprite Stage

The Voxa GridStage Sprite Stage is used with sections collected on to metal grids. A 3D rendering of the Sprite Stage is shown in Figure S1. It combines two linear piezo stages to have x and y translation. The axes of the Sprite Stage have a scan resolution of approximately 1 nm with repeatability of approximately 50 nm. The system was designed to translate Grid Sticks, which are cartridges that can hold up to 16 standard 3 mm TEM grids. Since Grid Sticks are much larger and more robust than individual TEM grids, sample handling errors were reduced while increasing the load density. An additional advantage is that Grid Sticks are a storage medium for standard sample grids, allowing for easy indexing when archiving thousands of sample grids.

### VOXA GridStage Reel

The Voxa GridStage Reel is an in-column sample conveyer system designed to handle TEM tissues on tape, enabling sample indexing with a barcode ID reader. Two reel housings are mounted on opposite sides of the TEM column. The vacuum load lock for inserting the GridStage sits on top of the feed reel. A tension sensor is located inside the load lock and the system uses the tension reading to automatically adjust the tape movement when changing apertures. The tape goes through a pinch-roller, wraps through the tension sensor, and then is fed into the channel of the GridStage. The tape is always tensioned during movement and is slack during montaging. Clamps at both ends of the GridStage ensure that the tape is fixed down onto the GridStage, eliminating tape slippage. Two additional camera viewports, at the feed and take-up reel housings, are used to monitor tape status during GridTape translation.

### autoTEM Column Extension

The autoTEM column extension continues the vacuum column of the microscope and is terminated with a custom scintillator coated with P43 phosphor, followed by a custom leaded glass window that blocks X-rays from escaping through the bottom of the column. The rest of the column extension is encased in lead shielding panels to block x-rays around the column. Below the column is a custom camera housing with a single camera (XIMEA CMOSIS CMV20000 or CMV50000) that images the scintillator screen. The PCI-E interface offered by the CMOSIS camera also provided sufficiently fast data transfer rates to enable on-the-fly GPU-based image processing and quality control.

### pyTEM Server

The pyTEM server runs on a Virtual Machine in the Allen Institute data center. The server hosts the pyTEM GUI web pages and provides a link between pyTEM and multiple web clients. It republishes the images and status streams coming from pyTEM to an arbitrary number of clients and acts as a gatekeeper, preventing contention between clients by only allowing a single client to issue commands. A control token can be grabbed by a client to take ownership of an EM, each of which is controlled by a single, dedicated server.

### pyTEM Function Highlights

#### TAO database and Automatic ROI Generation

Given that tissue sections can number in the tens of thousands, mapping tissue ROI becomes a labor-intensive task and requires an automation process as well. Therefore, TAO (TEM Acquisition Objects) are introduced by referencing ROIs through high-mag coordinate space only. TAO images are created from the images captured from the ATUM system during sectioning. The TAOUpload system creates these TAO objects and uploads them to AWS. This allows multiple users to simultaneously access this data, define ROIs, and perform QC verification of correct ROI definition.

We have also developed automated scripts to create tissue ROIs, aligning the EM imaging region to the functional imaging area. The tissue area on each aperture can be detected automatically from optical sectioning images (Figure S8). Based on pre-defined rules of absolute distance to the tissue corner along with tissue compression scaling factors, we achieved ~97% success rate of placing correct ROI’s. Once data is uploaded to AWS it is immediately available for use in TEM imaging or QC analysis. We have verified through 2D montage that the ROI placement is very precise with this method.

The automatic ROI generation sequence followed per aperture at present is as follows: (1) extract the barcode, (2) threshold the image to find the aperture candidates, (3) select the largest aperture candidate which is nearest the centerline of the optical image, (4) from the aperture extract the centroid and bounding box, (5) find the tissue area using HSV color segmentation over a range of colors, (6) filter out noise in the tissue area by removing small interior blobs and trimming small outer tendrils, (7) reject tissue candidates which don’t meet a minimum size criteria and extract the centroid of the tissue area, (8) optionally create a fixed size ROI at the tissue centroid, (9) insert all extracted data in to the TAO, and upload and the original image to AWS using the specimen_id, media_id, and barcode as keys for retrieval. If suitable tissue cannot be automatically located, an empty TAO placeholder is created and uploaded for eventual manual ROI definition.

In addition to the automatic ROI generation described above, pyTEM GUI is also a multi-resolution, web-based CAD system for manual ROI definition and editing at either low or high mag EM images or on optical images (Figure S3). At hi-mag it is possible to automatically locate the ROI corners to refine ROI placement, which may be required to correct for beam hysteresis when switching magnifications. When montaging, the user specifies a range of ROIs to image using a format similar to Microsoft Word page print selection (1-8, 13, 783-799).

#### Calibration

The nature of an electron beam is dependent upon the characteristics of the electron-generating filament and its alignment within the system. Thus, after initial installation and whenever a TEM filament is changed, the imaging subcomponents of the system need to be recalibrated. The most critical features to tune are the pixel resolution (nm/pixel) and the angular beam rotation. As part of the real-time QC module, the pixel calibration is done through template matching. The image pixel displacements of a 3×3 grid of template located at the center of the screen are measured for known stage displacements. The standard deviation of each X and Y for each vector is less than one pixel and results are now highly repeatable. Overall, this method ensures calibration consistency across multiple autoTEM systems.

#### Centroid Finding

The coordinate system used to place ROIs on the optical ATUM image assigns the aperture centroid as the (0, 0) reference point. ROIs are then defined in physical distance units as X/Y offsets from this centroid point along with a rotation angle. When an aperture is imaged, the tape subsystem first positions the aperture in the approximate center of the column. Next, the centroid of the aperture is detected in stage coordinates using a binary search algorithm for the light-dark transition point demarking each of the four aperture edges: top, bottom, left, right. Each ROI is then translated to this stage coordinate centroid, and further rotated by a magnification-specific beam rotation which is measured during the calibration process.

#### Brightfield Correction

To compensate for camera, lens, and illumination non-linearities, each montage image is corrected using previously acquired brightfield and darkfield images. The darkfield corrects sensor per-pixel dark current offsets and is acquired once per week, or whenever the camera gain is altered. The brightfield (example in Figure S9) corrects illumination gradients (which can change with electron beam drift), camera lens non-linearities, and sensor per-pixel gain variations. A new brightfield is acquired for each aperture.

The darkfield is acquired by averaging together 16 images with the EM beam turned off. The brightfield is acquired over tissue with the electron beam on while moving the stage slowly across a 300 μm radius region, integrating light from light and dark tissue regions. Sixteen such integrated images are averaged and then scaled to create the brightfield.

For each montage image, a corrected image is created using the brightfield and darkfield images on the GPU in floating point format:

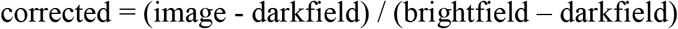

The corrected image is converted to 8bpp and saved as a TIFF file.

#### Auto Focus

Over the course of scaling montage area from (100 μm)^2^ to 1 mm^2^, the autofocus algorithm had to evolve to deal with a property of the JEOL 1200EXII TEM where overall image intensity changes with focus value. The current focus measurement is derived as follows (Figure S10): (1) from the corrected 3840 × 3840 image, extract the center 2048 × 2048 pixels, (2) compute the log magnitude of the Discrete Fourier Transform (DFT):

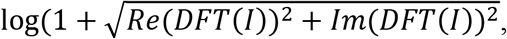

(3) create an image mask which excludes lowest 6 frequency components and high frequency components above 1600, (4) perform a polar to rectangular transformation of the DFT components within the mask area, (5) find the average of the components for each row, (6) sum the averages. Discarding the highest frequency components (which are largely noise) makes the algorithm less susceptible to the image intensity changes that happen with the change of focus. Determination of optimal FFT frequency components was derived experimentally by comparing FFT scores at different exposures and different frequency components.

Through the experiments from pilot datasets, we noticed that a single EM focus point at the ROI centroid was not sufficient for a large 1 mm^2^ montage. We sometimes failed to derive optimal focal point when a blood vessel or other void existed at the ROI centroid; or when there was a sample height gradient across ROI due to tissue section being placed towards the aperture edge. To improve the autofocus algorithm, pyTEM now performs focus optimization at additional satellite points distributed across the ROI, averaging the remaining measurement points to create a reasonably optimal focus value for the whole montage.

#### Performance tuning (LaB6, Zeiss Lens, stage performance)

To achieve the optimal speed and quality of the image platform, we have also made improvements in terms of electron source, camera lens and stage performance. The electron source for the autoTEM system is a critical component for long term imaging experiments. Traditionally the JEOL 1200EXII leverages both Tungsten hairpin style filaments and Lanthanum hexaboride crystal (LaB_6_) as possible electron sources. Tungsten filaments, although easier to operate, demonstrated poor stability for long periods of time due to deterioration of the tungsten source, as well as a short life span (~100 hours). LaB6 filaments have a more stringent ultimate vacuum pressure but have a much larger current density (higher electron flux per unit area) which reduces exposure times for the capture camera as well as a much longer lifetime (1,000-2,000 hrs). The average lifetime by using LaB6 filaments during production imaging was about 1 month for continuous 24/7 operation. In addition, the exposure time using the tungsten source was 100-150 ms as compared to 50-80 ms using LaB6, with same camera, optical arrangement and specimen. Among the 300 ms acquisition cycle for each tile, the stage "step-and-settle” time from initializing the stage move to finishing the move and send the “complete” status back to pyTEM is averaged at ~120 ms, consuming almost half of the total cycle time. Therefore, it is very important to optimize the stage performance parameters. We focused on SmarAct stage speed, acceleration and dwell time for stage to settle down. A combination of various stage speeds and accelerations were tested. In general, we have found a linear correlation between SmarAct stage acceleration and speed to the cycle time: the larger the acceleration, the less the “step-and-settle” time; the higher the speed, the less the “step-and-settle” time but caps around 10 mm/s. An appropriate dwell time is also necessary, otherwise the random streaks of blurry tiles appear on the montage, because of insufficient wait time for stage to settle and camera exposure before the next move. Larger frame size also requires more dwell time because of increasing step distance for each tile. We obtained “step-and-settle” time at 120 ms by average among six stages for the current setup.

#### Tissue Preparation

All procedures were carried out in accordance with the Institutional Animal Care and Use Committee at the Allen Institute for Brain Science. Mice were transcardially perfused with a fixative mixture of 2.5% paraformaldehyde and 1.25% glutaraldehyde. After dissection, slices were cut with a vibratome and post-fixed for 12 – 48 h. Slices were extensively washed and prepared for reduced osmium treatment (rOTO) based on the protocol of Hua and colleagues^27^. Ferricyanide was used to reduce Osmium and Thiocarbohydrazide (TCH) for further intensification of the staining. Uranyl acetate and lead aspartate were used to enhance contrast. After resin embedding, ultrathin sections (40 nm) were either manually cut in a Leica ultra-microtome our automatically onto GridTape using an RMC Automated Tape Collecting Ultramicrotome.

#### Sharing of software and hardware design

We will share software, bill of materials, and hardware design drawings. Software will be shared under the Allen Institute Software License and Contribution Agreement, subject to any applicable third-party licensing restrictions. Bill of materials and hardware design will be shared under the Allen Institute Terms of Use: https://alleninstitute.org/legal/terms-use/.

## Supplemental Information

**Figure S1:**
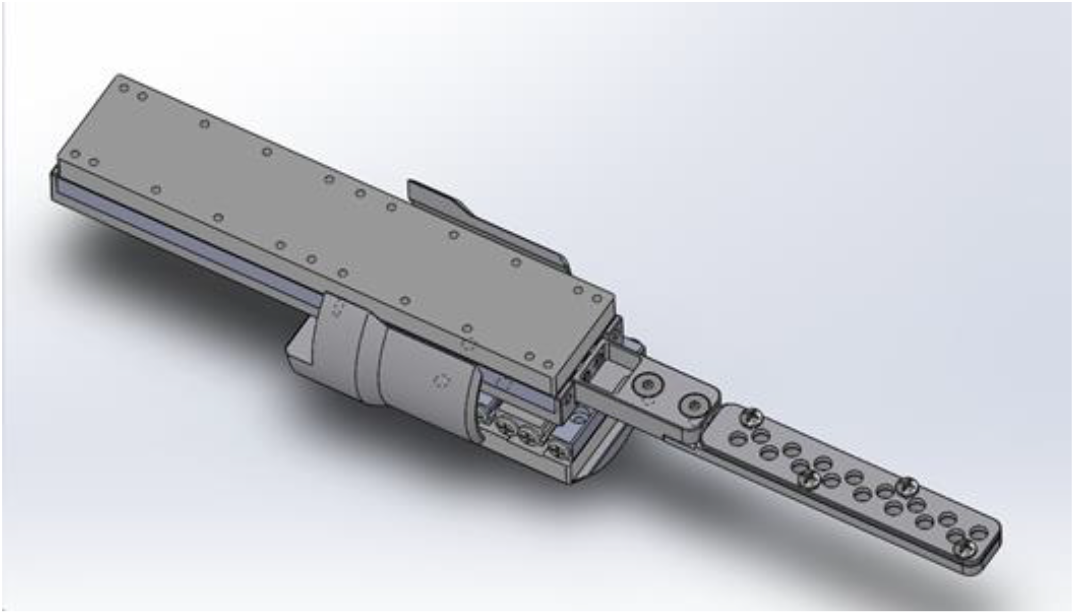
VOXA GridStage Sprite Stage. A 3D rendering of the Sprite Stage used to image sections collected onto standard grids. Note the Grid Stick has 16 wells for holding standard TEM grids.

**Figure S2.**
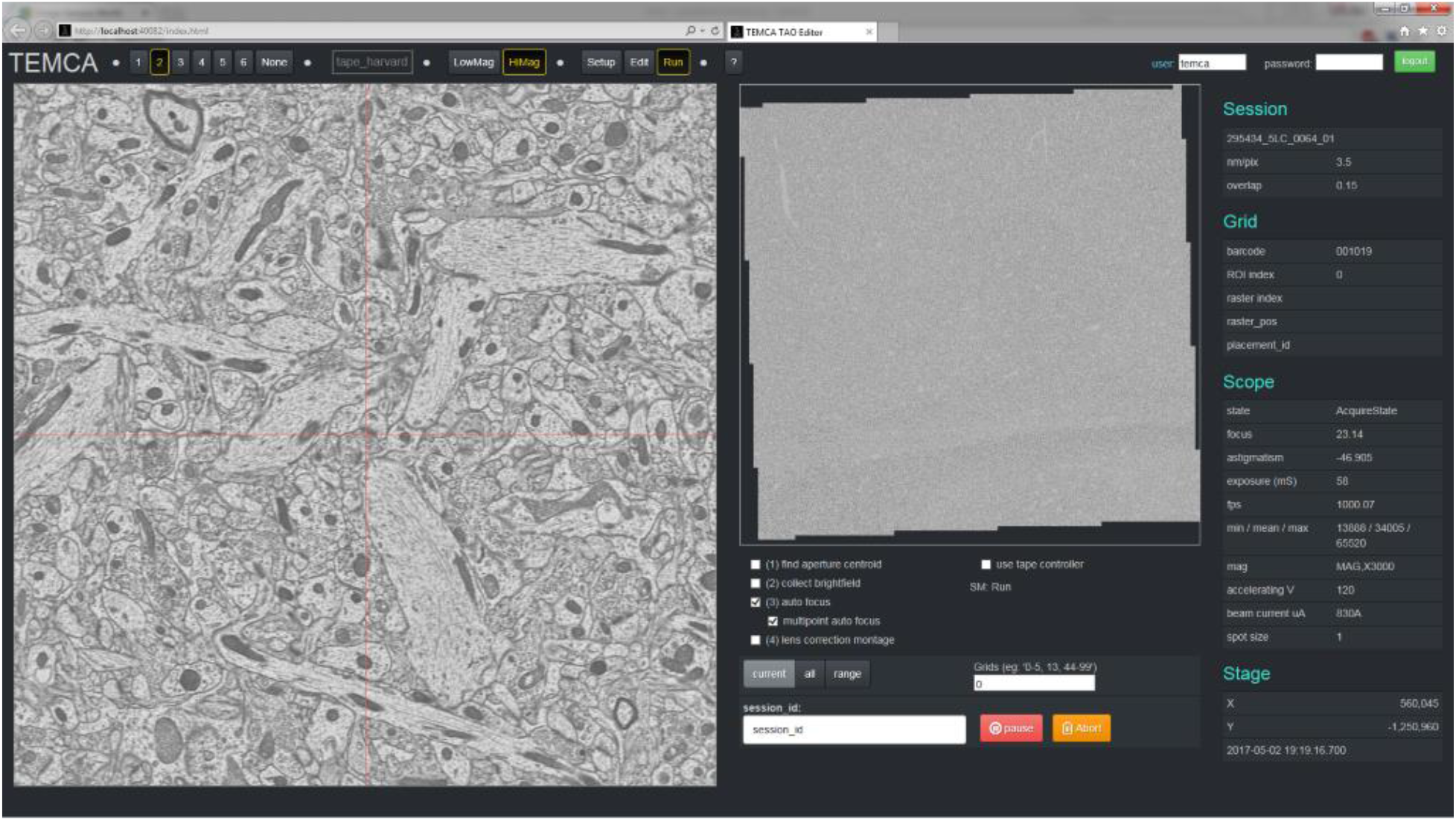
pyTEM GUI. The pyTEM GUI provides the user with a web-based interface to perform manual imaging as well as set up long imaging runs. The right panel displays the real-time running parameters such as the section information, scope and stage status.

**Figure S3.**
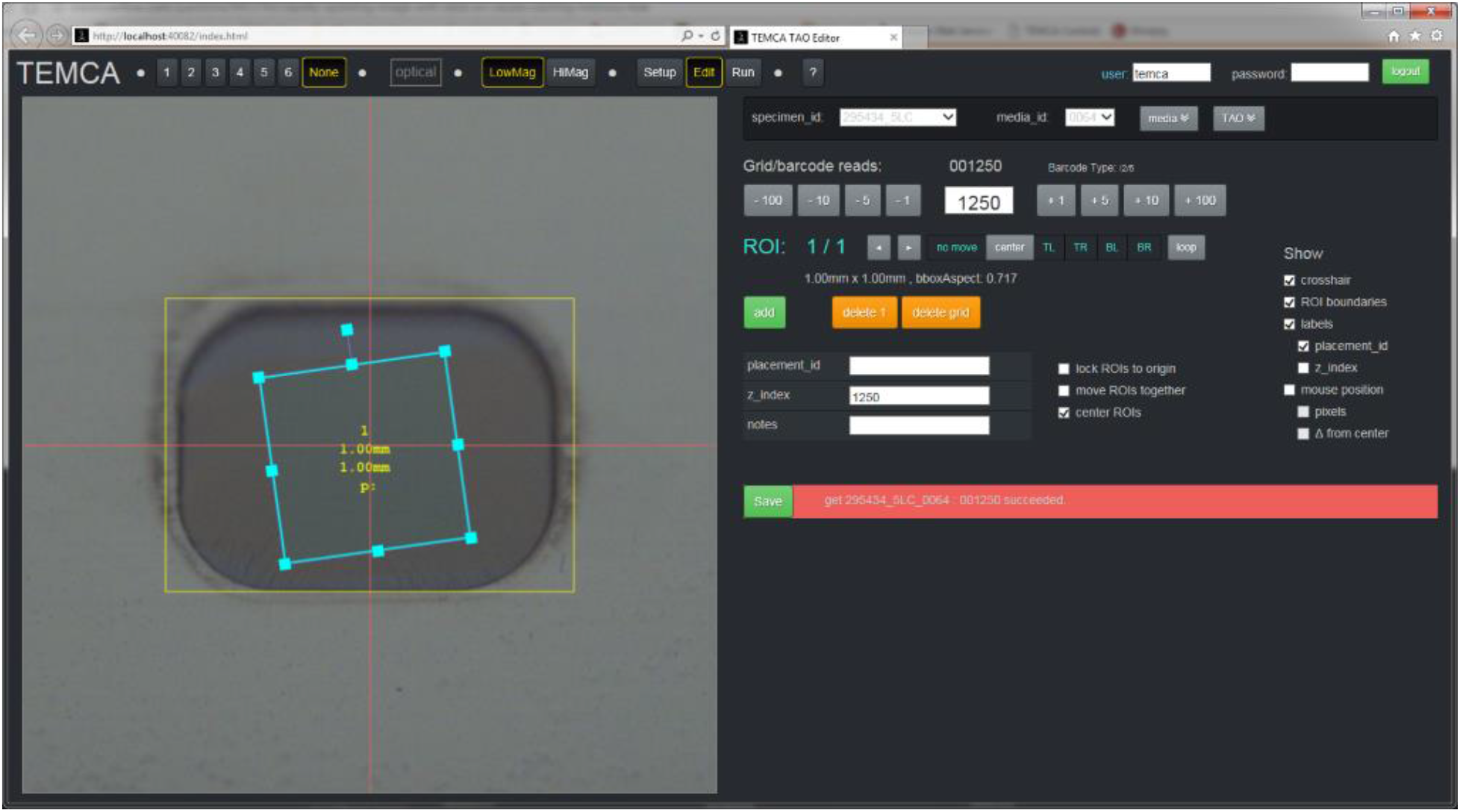
User interface to define ROIs for high-fidelity raster imaging with pyTEM GUI.

**Figure S4:**
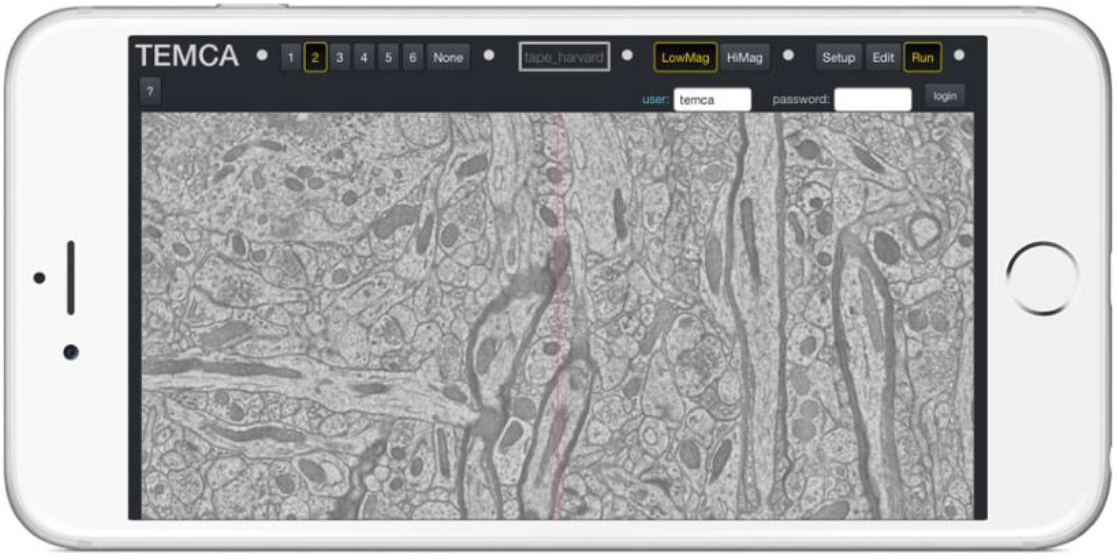
pyTEM GUI utilizes a web-based interface that allows supervision and control of autoTEM operation from multiple clients and using any web-connected device with requisite security credentials.

**Figure S5:**
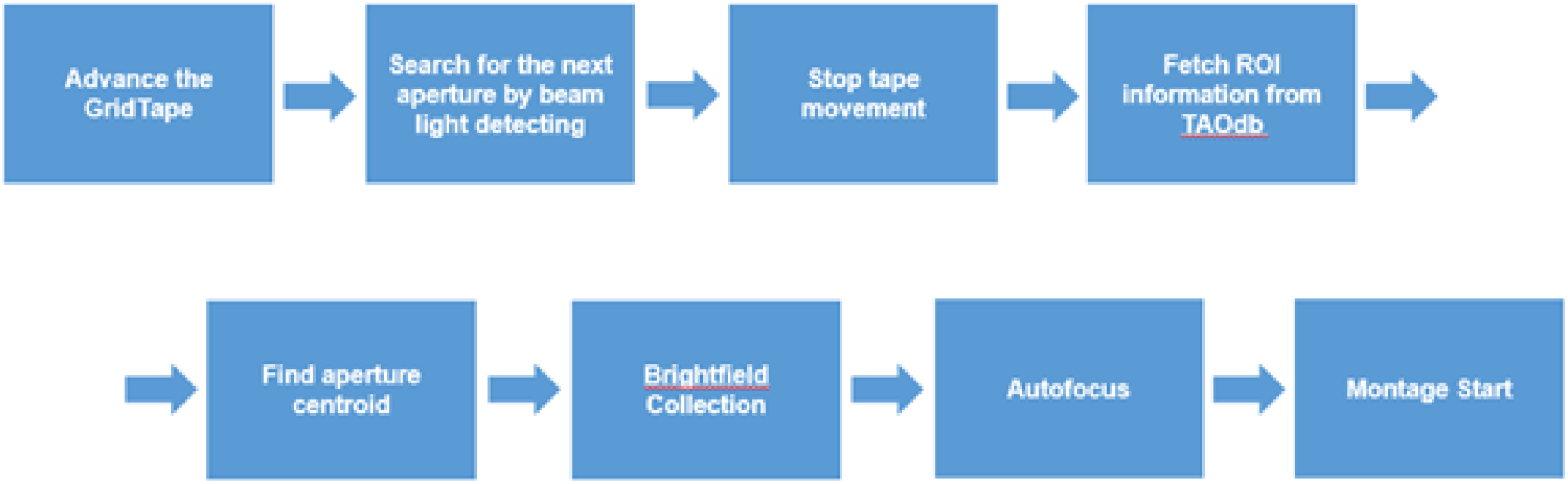
Automated Flow for montage acquisition. The tape is advanced to the desired ROI. An ROI associated with the aperture is pulled from AWS. The center of the aperture is found via edge detection. A brightfield is collected and applied to all subsequent images linked with the ROI. Finally, an autofocus operation is performed to maximize image quality and then the raster scan is initiated.

### Multi-System Monitoring (MSM)

With increasing demands for imaging larger tissue volumes, high TEM system availability and minimized downtime are required for continuous imaging. Traditionally most automated electron microscopy projects are centered on one imaging machine whereas the piTEAM pipeline requires multiple machines running in parallel. The need to quickly diagnose failures in a multi-system environment requires a robust method of troubleshooting as well as active preventative maintenance. In order to satisfy these requirements, a monitoring system for piTEAM and the laboratory environment was implemented.

The autoTEM Multi System Monitoring (MSM) is a combination of a server and web-based UI (Grafana, Grafana Labs) that allows the users to quickly ascertain if any subcomponent needs attention. If a subcomponent is detected to have an issue, MSM communicates with pyTEM and informs the appropriate individual(s) that an error has occurred. In catastrophic cases, MSM can also safely pause the acquisition of an autoTEM system to prevent system damage and potential sample loss. The system was designed to track the status of any subcomponent issue and communicate alerts to users via automated message communication protocols (email, SMS, or Slack).

MSM tracks three top level systems that each contain an assembly of sub-systems: Facilities, Environment, and Equipment. Facility related sub-systems include the status of the pressure, flow, and temperature of chilled water circulating within each system. The environmental sub-systems include the status of laboratory conditions such as room temperature and humidity. Equipment sub-systems include the status of each autoTEM including filament life, vacuum pressure, and beam current. Image acquisition parameters are also tracked including focus scores, current machine activity, CPU, and memory use, and local disk space availability.

**Figure S6:**
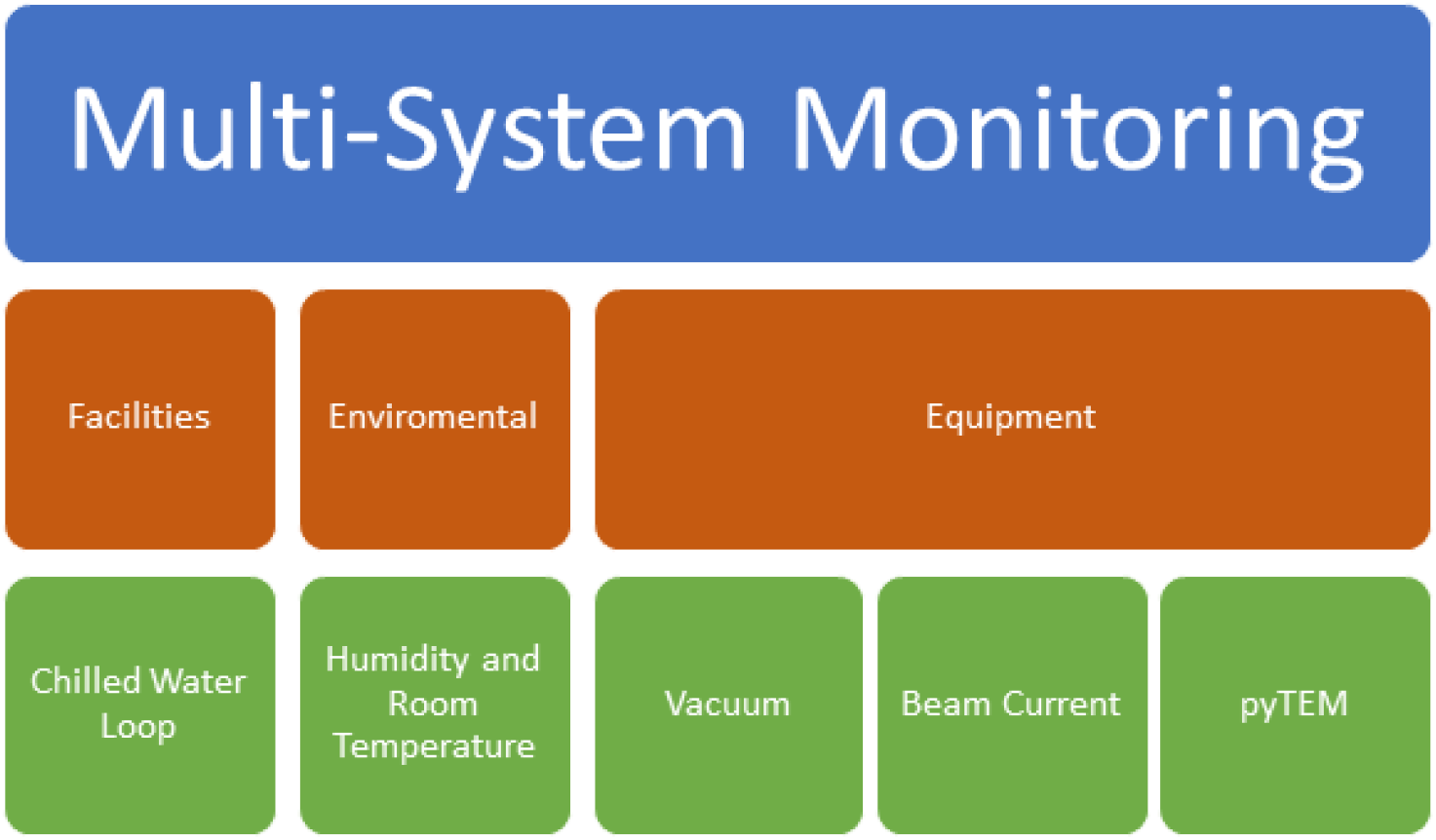
Hierarchical schematic of the autoTEM Multi System Monitoring. Bottom up reporting of sub-components and sensors are integrated into a unified monitoring software system for easy diagnosing. All information is logged to an AWS database.

**Figure S7:**
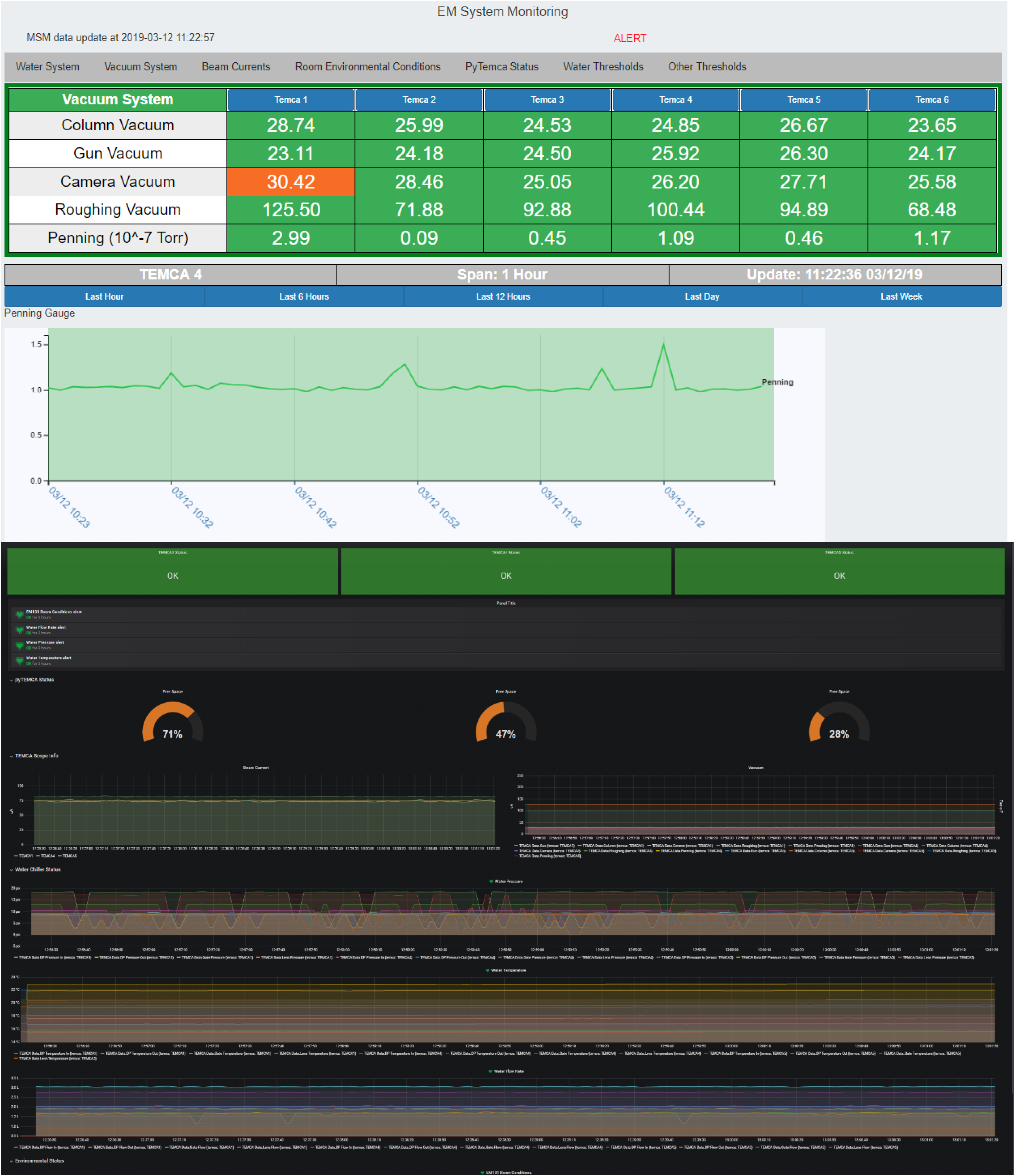
Screenshot of the MSM GUI monitoring the vacuum levels of the autoTEMs.

**Figure S8:**
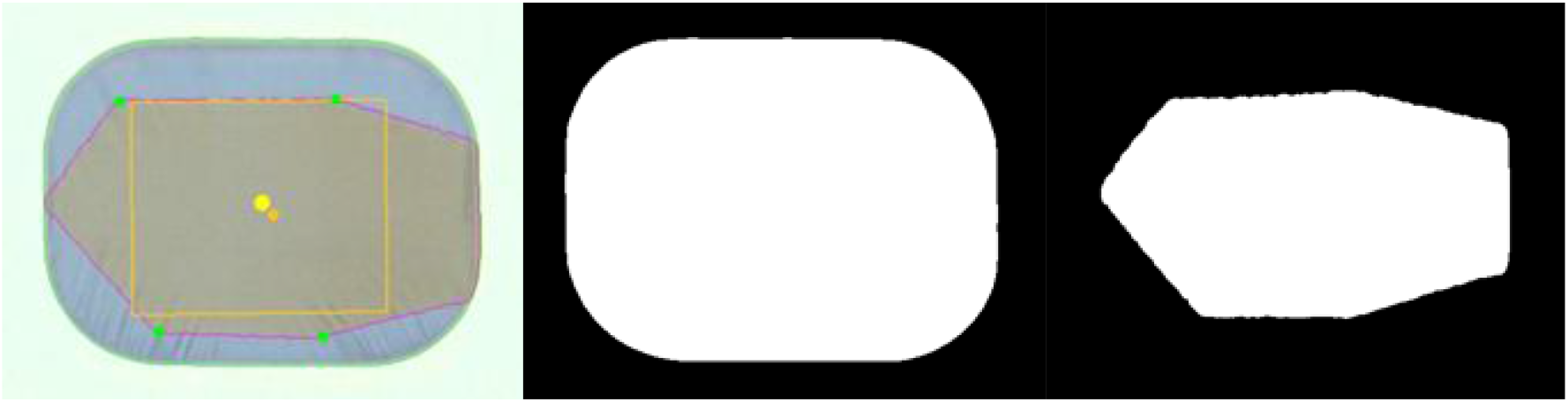
Auto ROI definition from an optical aperture image. The aperture and tissue boundaries are automatically detected, and an ROI is placed according to the distance offset and tissue compression scaling factor.

**Figure S9:**
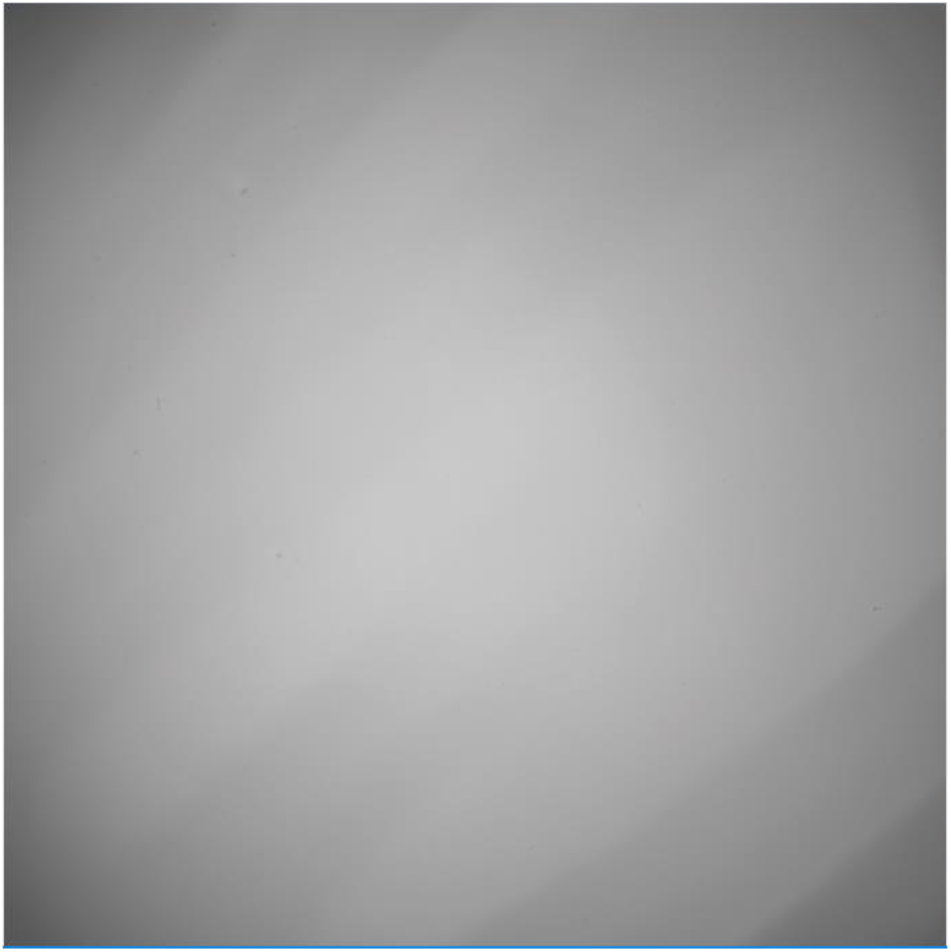
Example brightfield collected from an autoTEM using 50Mpix camera.

**Figure S10.**
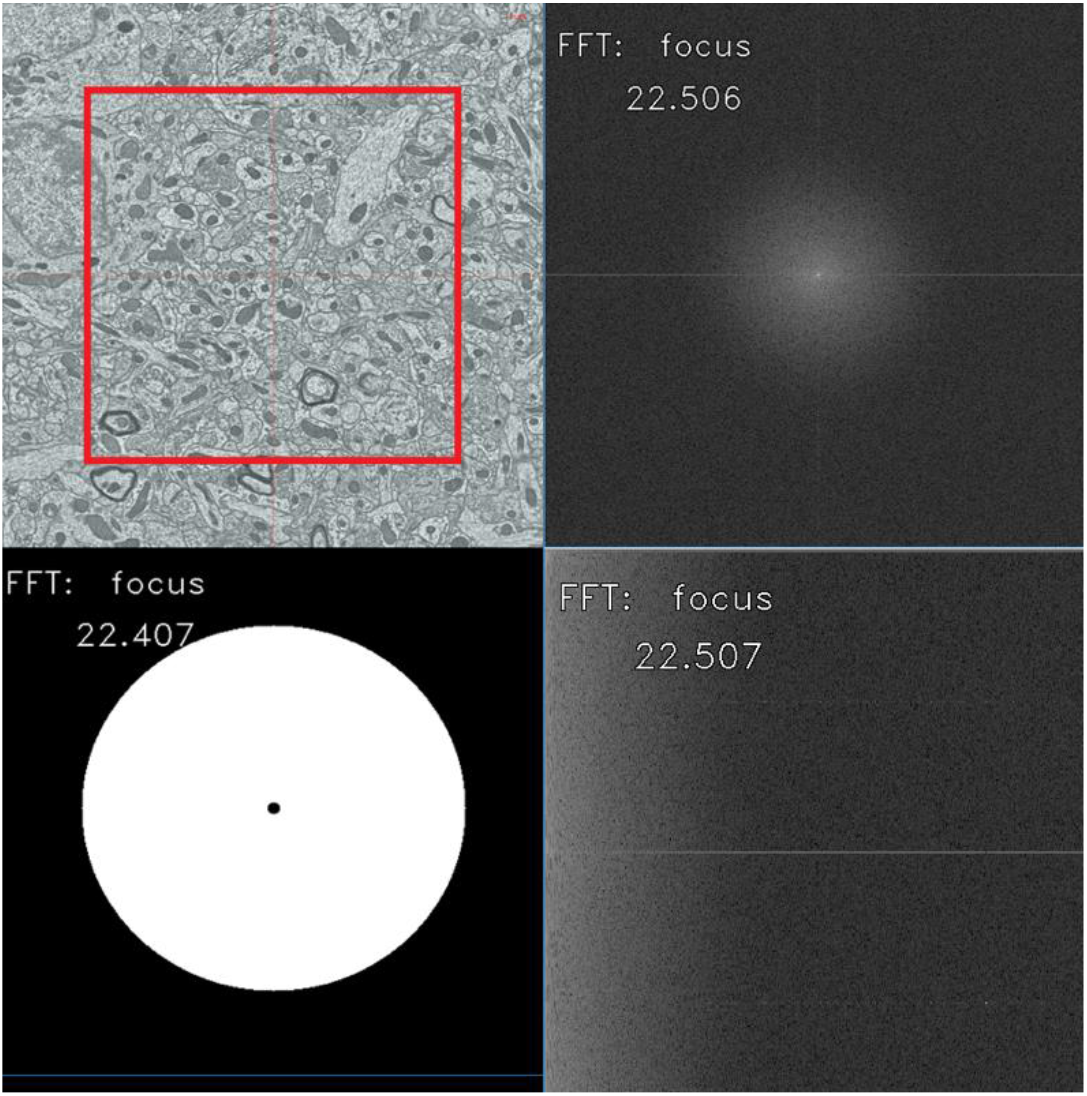
TL: center 2048×2048 image from which focus is to be measured, TR: DFT, BL: frequency range mask, BR: polar to rectangular conversion of DFT within mask.

### Montage QC Failure Examples

Figure S11 shows an example for tape slippage, in which case the GridTape is not secured in-place on the stage and shifts the position during stage movement. As a result, the montage quality map highlights misaligned rows and areas, or sometimes seen as partially imaged ROI. During 1 mm^3^ imaging, most of the reimaging needs came from tile overlap issue. Due to stage movement errors, there was a small fraction of tiles (< 1%) that did not have enough overlap to their neighboring tiles, usually along x-axis, and thus the image processing pipeline had difficulty drawing point matchers from the overlap region (Fig. S11b) to perform the stitching and alignment. We have also seen “random blurry tiles” (Fig. S11c) occurring because of charging, autofocus failure, or the stage not having enough settling time and thus a frame is captured by camera while stage is still moving. As an example, focus map in (Fig. S11d) shows a gradient across the montage where the right side is impacted by electron charging and tiles are slightly blurry.

**Figure S11:**
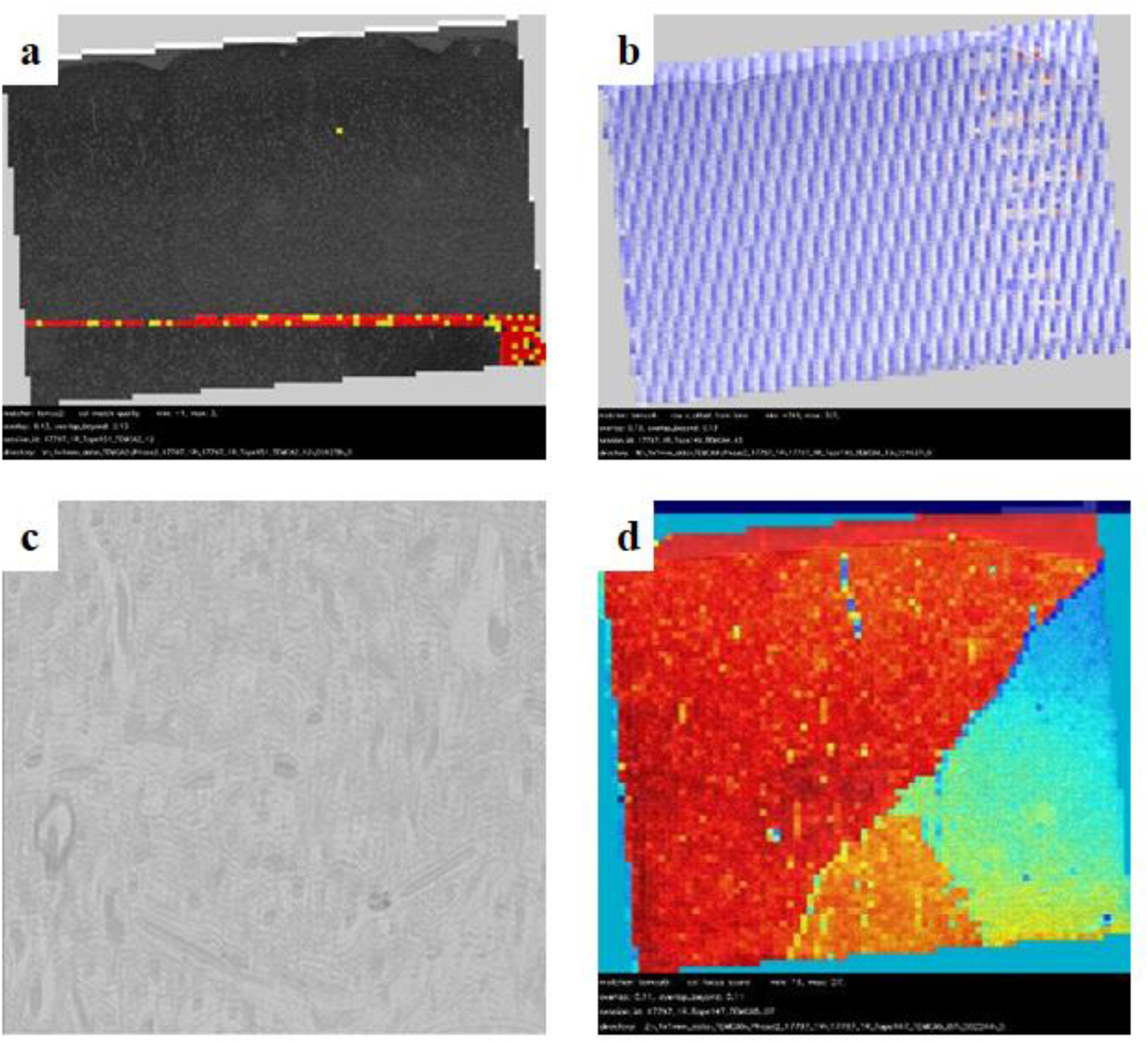
a. Tape slippage detected as flagged rows; b. Insufficient overlap seen as non-uniform pattern. c. Random blurry tile; d. Focus gradient due to charging.

**Figure S12:**
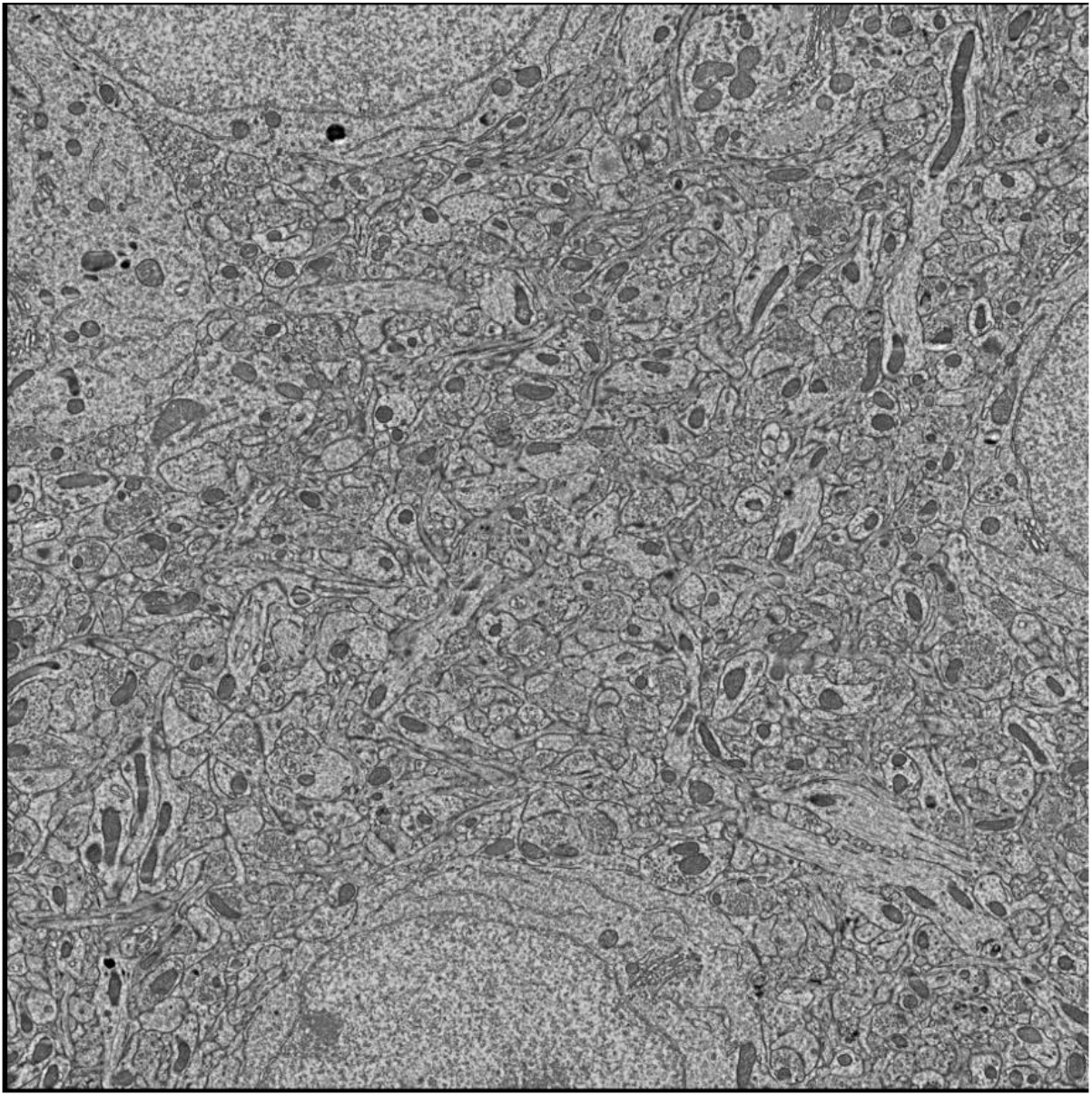
High-res image collected by AMT lens assembly using a 50MP XIMEA camera. The scope magnification is at 2500x. The pixel resolution is approximately 4nm/pixel.

**Table S13:** Imaging metric comparison between current 50MP sensor vs future 100MP sensor (projected). The values are calculated using 4nm / pixel spatial resolution. The ROI size is 1 mm2.

| Camera Model | XiB 50 Mpixel |  | > 100 Mpixel |  |
| --- | --- | --- | --- | --- |
|  | X | Y | X | Y |
| **Cropped Frame (pixels)** | 5408 | 5408 | 10000 | 10000 |
| **FOV (microns)** | 22 | 22 | 40 | 40 |
| **Overlap %** | 9 | 9 | 5 | 5 |
| **Tiles w/overlap** | 54 | 54 | 28 | 28 |
| **Total Tiles/Section** | 2916 |  | 784 |  |

